# Organohalide-respiring *Desulfoluna* species isolated from marine environments

**DOI:** 10.1101/630186

**Authors:** Peng Peng, Tobias Goris, Yue Lu, Bart Nijsse, Anna Burrichter, David Schleheck, Jasper J. Koehorst, Jie Liu, Detmer Sipkema, Jaap S. Sinninghe Damste, Alfons J. M. Stams, Max M. Häggblom, Hauke Smidt, Siavash Atashgahi

**Affiliations:** Laboratory of Microbiology, Wageningen University & Research, Stippeneng 4, 6708 WE Wageningen, The Netherlands; Department of Applied and Ecological Microbiology, Institute of Microbiology, Friedrich Schiller University, 07743 Jena, Germany; College of Environmental Science and Engineering, Hunan University, 410082 Changsha, China; Laboratory of Systems and Synthetic Biology, Wageningen University & Research, Stippeneng 4, 6708 WE Wageningen, The Netherlands; Department of Marine Microbiology and Biogeochemistry, NIOZ Royal Netherlands Institute for Sea Research, P.O. Box 59, 1790 AB Den Burg, The Netherlands; Department of Earth Sciences, Faculty of Geosciences, Utrecht University, P.O. Box 80.121, 3508 TA Utrecht, The Netherlands; Centre of Biological Engineering, University of Minho, Campus de Gualtar, 4710-057 Braga, Portugal; Department of Biochemistry and Microbiology, Rutgers University, New Brunswick, NJ 08901, USA; Department of Biology, University of Konstanz, 78457 Konstanz, Germany; The Konstanz Research School Chemical Biology, University of Konstanz, 78457 Konstanz, Germany

**Author notes:** Address correspondence to Siavash Atashgahi.

## Abstract

The genus *Desulfoluna* comprises two anaerobic sulfate-reducing strains, *D. spongiiphila* AA1^⊤^ and *D. butyratoxydans* MSL71^⊤^ of which only the former was shown to perform organohalide respiration (OHR). Here we isolated a third member of this genus from marine intertidal sediment, designed *D. spongiiphila* strain DBB. All three *Desulfoluna* strains harbour three reductive dehalogenase gene clusters (*rdhABC*) and corrinoid biosynthesis genes in their genomes. Brominated but not chlorinated aromatic compounds were dehalogenated by all three strains. The *Desulfoluna* strains maintained OHR in the presence of 20 mM sulfate or 20 mM sulfide, which often negatively affect OHR. Strain DBB sustained OHR with 2% oxygen in the gas phase, in line with its genetic potential for reactive oxygen species detoxification. Reverse transcription-quantitative PCR (RT-qPCR) revealed differential induction of *rdhA* genes in strain DBB in response to 1,4-dibromobenzene or 2,6-dibromophenol. Proteomic analysis confirmed differential expression of *rdhA1* with 1,4-dibromobenzene, and revealed a possible electron transport chain from lactate dehydrogenases and pyruvate oxidoreductase to RdhA1 via menaquinones and either RdhC, or Fix complex (electron transfer flavoproteins), or Qrc complex (Type-1 cytochrome c3:menaquinone oxidoreductase).

## Introduction

More than 5,000 naturally produced organohalides have been identified, some of which have already been present in a variety of environments for millions of years [1]. In particular, marine environments are a rich source of chlorinated, brominated and iodinated organohalides produced by marine algae, seaweeds, sponges, and bacteria [2], Fenton-like [3] and photochemical reactions, as well as volcanic activities [4, 5]. Such a natural and ancient presence of organohalogens in marine environments may have primed development of various microbial dehalogenation metabolisms [6]. Furthermore, marine environments and coastal regions in particular are also commonly reported to be contaminated with organohalogens from anthropogenic sources [7].

During organohalide respiration (OHR) organohalogens are used as electron acceptors, and their reductive dehalogenation is coupled to energy conservation [8–10]. This process is mediated by reductive dehalogenases (RDases), which are membrane-associated, corrinoid-dependent, and oxygen sensitive proteins [9–11]. The corresponding *rdh* gene clusters usually consists of *rdhA* encoding the catalytic subunit, *rdhB* encoding a putative membrane anchor protein [10], and a variable set of accessory genes encoding RdhC and other proteins likely involved in regulation, maturation and/or electron transport [12, 13]. The electron transport chain from electron donors to RDases has been classified into quinone-dependent (that rely on menaquinones as electron shuttles between electron donors and RDases) and quinone-independent pathways [9, 10, 14]. Recent studies suggested that RdhC may serve as electron carrier during OHR in *Firmicutes* [15, 16].

OHR is mediated by organohalide-respiring bacteria (OHRB), which belong to a broad range of phylogenetically distinct bacterial genera. OHRB belonging to *Chloroflexi* and the genus *Dehalobacter* (*Firmicutes*, e.g. *Dehalobacter restrictus*) are specialists restricted to OHR, whereas proteobacterial OHRB and members of the genus *Desulfitobacterium* (*Firmicutes*, e.g. *Desulfitobacterium hafniense)* are generalists with a versatile metabolism [17, 18]. Numerous studies have reported OHR activity and occurrence of OHRB and *rdhA* genes in marine environments [6, 19–21]. Recent genomic [22–24] and single-cell genomic [25] analyses revealed widespread occurrence of *rdh* gene clusters in marine *Deltaproteobacteria*, indicting untapped potential for OHR. Accordingly, OHR metabolism was experimentally verified in three *Deltaproteobacteria* strains, not previously known as OHRB [23].

OHRB, and in particular members of the *Chloroflexi*, are fastidious microbes, and are susceptible to inhibition by oxygen [26], sulfate [27] or sulfide [28, 29]. In the presence of both 3-chlorobenzoate and either sulfate, sulfite or thiosulfate, *Desulfomonile tiedjei* isolated from sewage sludge preferentially performed sulfur oxyanion reduction [30], and OHR inhibition was suggested to be caused by downregulation of *rdh* gene expression [30]. In contrast, concurrent sulfate reduction and OHR was observed in *Desulfoluna spongiiphila* AA1^⊤^ isolated from the marine sponge *Aplysina aerophoba* [20], and three newly characterized organohalide-respiring marine deltaproteobacterial strains [23]. Sulfate- and sulfide-rich marine environments may have exerted a selective pressure resulting in development of sulfate- and sulfide-tolerant OHRB.

The genus *Desulfoluna* comprises two anaerobic sulfate-reducing strains, *D. spongiiphila* AA1^⊤^ isolated from the bromophenol-producing marine sponge *Aplysina aerophoba* [20, 31], and *D. butyratoxydans* MSL71^⊤^ isolated from estuarine sediments [32]. Strain AA1^⊤^ can reductively dehalogenate various bromophenols but not chlorophenols. The genome of strain AA1^⊤^ harbours three *rdhA* genes, one of which was shown to be induced by 2,6-dibromophenol [21]. The OHR potential and the genome of strain MSL71^⊤^ have not been studied before. In this study, a third member of the genus *Desulfoluna*, designated *D. spongiiphila* strain DBB, was isolated from a marine intertidal sediment. The OHR metabolism of strain DBB and of strain MSL71^⊤^ was verified in this study. In line with former reports [22–25], this study further reinforces an important role of marine organohalide-respiring *Deltaproteobacteria* in halogen, sulfur and carbon cycling.

## Materials and Methods

### Chemicals

Brominated, iodinated and chlorinated benzenes and phenols were purchased from Sigma-Aldrich. Other organic and inorganic chemicals used in this study were of analytical grade.

### Bacterial strains

*D. spongiiphila* AAΓ (DSM 17682^⊤^) and *D. butyratoxydans* MSL71^⊤^ (DSM 19427^⊤^) were obtained from the German Collection of Microorganisms and Cell Cultures (DSMZ, Braunschweig, Germany), and were cultivated as described previously [20, 32].

### Enrichment, isolation and cultivation of strain DBB

Surface sediment of an intertidal zone, predominantly composed of shore sediment, was collected at the shore in L’Escala, Spain (42°7’35.27”N 3°8’6.99”E). Five grams of sediment were transferred into 120-ml bottles containing 50 ml of anoxic medium [33] with lactate and 1,4-dibromobenzene (1,4-DBB) as the electron donor and acceptor, respectively. Sediment-free cultures were obtained by transferring the suspensions of the enrichment culture to fresh medium. A pure culture of a 1,4-DBB debrominating strain, designated as *D. spongiiphila* strain DBB, was obtained from a dilution series on solid medium with 0.8% low-melting point agarose (Sigma-Aldrich). A detailed description of enrichment, isolation and physiological characterization of strain DBB is provided in the Supplementary Information.

### Cell morphology and cellular fatty acids analyses

Cell morphology and motility were observed using a LEICA DM 2000 Microscope and a JEOL-6480LV Scanning Electron Microscope (SEM). Actively growing cells were directly observed under the 100x magnification objective of the LEICA DM 2000 Microscope. Sample fixation and dehydration for SEM were performed as described previously [34]. The cellular fatty acid composition was analysed from 500 ml cultures of AA1^⊤^, DBB and MSL71^⊤^, which were grown with 20 mM lactate and 10 mM sulfate. Fatty acids in the cell were analysed by acid hydrolysis of total cell material following a method previously described [35].

### DNA extraction and bacterial community analysis

DNA of the intertidal sediment (5 g) and the 1,4-DBB-respiring enrichment culture (10 ml) was extracted using the DNeasy PowerSoil Kit (MO-BIO, CA, USA). A 2-step PCR strategy was applied to generate barcoded amplicons from the V1—V2 region of bacterial 16S rRNA genes as described previously [36]. Sequence analysis was performed using NG-Tax [37]. Operational taxonomic units (OTUs) were assigned taxonomy using uclust [38] in an open reference approach against the SILVA 16S rRNA gene reference database (LTPs128_SSU) [39]. Finally, a biological observation matrix (biom) file was generated and sequence data were further analyzed using Quantitative Insights Into Microbial Ecology (QIIME) v1.2 [40].

### Genome sequencing and annotation

DNA of DBB and MSL71^⊤^ cells was extracted using the MasterPure™ Gram Positive DNA Purification Kit (Epicentre, WI, USA). The genomes were sequenced using the Illumina HiSeq2000 paired-end sequencing platform (GATC Biotech, Konstanz, Germany). The genome of strain DBB was further sequenced by PacBio sequencing (PacBio RS) to obtain longer read lengths. Optimal assembly kmer size for strain DBB was detected using kmergenie (v. 1.7039) [41]. A *de novo* assembly with Illumina HiSeq2000 paired-reads was made with assembler Ray (v2.3.1) [41] using a kmer size of 81. A hybrid assembly for strain DBB with both the PacBio and the Illumina HiSeq reads was performed with SPAdes (v3.7.1, kmer size: 81) [42]. The two assemblies were merged using the tool QuickMerge (v1) [43]. Duplicated scaffolds were identified with BLASTN [44] and removed from the assembly. Assembly polishing was performed with Pilon (v1.21) [45] using the Illumina HiSeq reads. Optimal assembly kmer size for strain MSL71^⊤^ was also identified using kmergenie (v.1.7039), and a *de novo* assembly with Illumina HiSeq2000 paired-end reads was performed with SPAdes (v3.11.1) with a kmer-size setting of 79,101,117. FastQC and Trimmomatic (v0.36) [46] was used for read inspection and trimming using the trimmomatic parameters: TRAILING:20 LEADING:20 SLIDINGWINDOW:4:20 MINLEN:50. Trimmed reads were mapped with Bowtie2 v2.3.3.1 [47]. Samtools (v1.3.1) [48] was used for converting the bowtie output to a sorted and indexed bam file. The assembly was polished with Pilon (v1.21).

### Transcriptional analysis of the *rdhA* genes of *D. spongiiphila* DBB

Transcriptional analysis was performed using DBB cells grown with lactate (20 mM), sulfate (10 mM) and either 1,4-DBB (1 mM) or 2,6-DBP (0.2 mM). DBB cells grown with lactate and sulfate but without any organohalogens were used as control. Ten replicate microcosms were prepared for each experimental condition, and at each sampling time point, two microcosms were randomly selected and sacrificed for RNA isolation as described previously [49]. RNA was purified using RNeasy columns (Qiagen, Venlo, The Netherlands) followed by DNase I (Roche, Almere, The Netherlands) treatment. cDNA was synthesized from 200 ng total RNA using superScript™ III Reverse Transcriptase (Invitrogen, CA, USA) following manufacturer’s instructions. RT-qPCR assays were performed as outlined in Supplementary Information.

### Protein extraction and proteomic analysis

Triplicate cultures of strain DBB grown with lactate/sulfate (LS condition) or lactate/sulfate/1,4-DBB (LSD condition) were used for proteomic analysis. Preparation of cell-free extracts (CFE), determination of protein concentration, SDS-PAGE purification of total proteins in CFE and of proteins in membrane fragments, and the peptide fingerprinting-mass spectrometry (PF-MS) analysis, were performed as outlined in Supplementary Information. Statistical analysis was performed using prostar proteomics [50]. Top three peptide area values were normalized against all columns. The values of proteins detected in at least two of the three replicates were differentially compared and tested for statistical significance. Missing values were imputed using the SLSA function of prostar, and hypothesis testing with a student’s t-test was performed for LSD vs LS growth conditions. The p-values were Benjamini-Hochberg corrected and proteins with p-values below 0.05 and a log2 value of 1 or larger were considered statistically significantly up- or downregulated.

### Analytical methods

Halogenated benzenes and benzene were analyzed on a GC equipped with an Rxi-5Sil capillary column (Retek, PA, USA) and a flame ionization detector (GC-FID, Shimadzu 2010). Halogenated phenols and phenol were analyzed on a Thermo Scientific Accela HPLC System equipped with an Agilent Poroshell 120 EC-C18 column and a UV/Vis detector. Organic acids and sugars were analyzed using a ThermoFisher Scientific SpectraSYSTEM™ HPLC equipped with an Agilent Metacarb 67H column and RI/UV detectors. Sulfate, sulfite and thiosulfate were analyzed using a ThermoFisher Scientific Dionex™ ICS-2100 Ion Chromatography System equipped with a Dionex Ionpac analytical column and a suppressed conductivity detector. Cell growth was determined by measuring OD_600_ using a WPA CO8000 cell density meter (Biochrom, Cambridge, UK). Sulfide was measured by a photometric method using methylene blue as described previously [51].

### Strain and data availability

*D. spongiiphila* strain DBB was deposited in DSMZ under accession number DSM 104433. The 16S rRNA gene sequences of strain DBB were deposited in GenBank (accession numbers: MK881098—MK881099). The genome sequences of strains DBB and MSL71 were deposited in the European Bioinformatics Institute (EBI, Project ID: PRJEB31368). A list of proteins detected from strain DBB under LS and LDS growth conditions is available in Supplementary Datasets 1 (Soluble fraction) and 2 (Membrane fraction).

## Results and discussion

### Enrichment of 1,4-DBB debrominating cultures and isolation of strain DBB

Reductive debromination of 1,4-DBB to bromobenzene (BB) and benzene was observed in the original cultures containing intertidal sediment (Fig. 1A, 1B). Debromination of 1,4-DBB was maintained in the subsequent sediment-free transfer cultures (Fig. 1C). However, benzene was no longer detected and BB was the only debromination product, indicating loss of the BB-debrominating population. Up to date, the only known OHRB that can debrominate BB to benzene is *Dehalococcoides mccartyi* strain CBDB1 [52]. 1,4-DBB debromination to BB was stably maintained during subsequent transfers (data not shown) and after serial dilution (Fig. 1D). Bacterial community analysis showed an increase in the relative abundance of *Deltaproteobacteria* from ~2% in the intertidal sediment at time zero to ~13% after 104 days of enrichment (Fig. 1E). The genus *Desulfoluna* was highly enriched and comprised more than 80% relative abundance in the most diluted culture (10^7^ dilution) (Fig. 1E).

**Figure 1.**
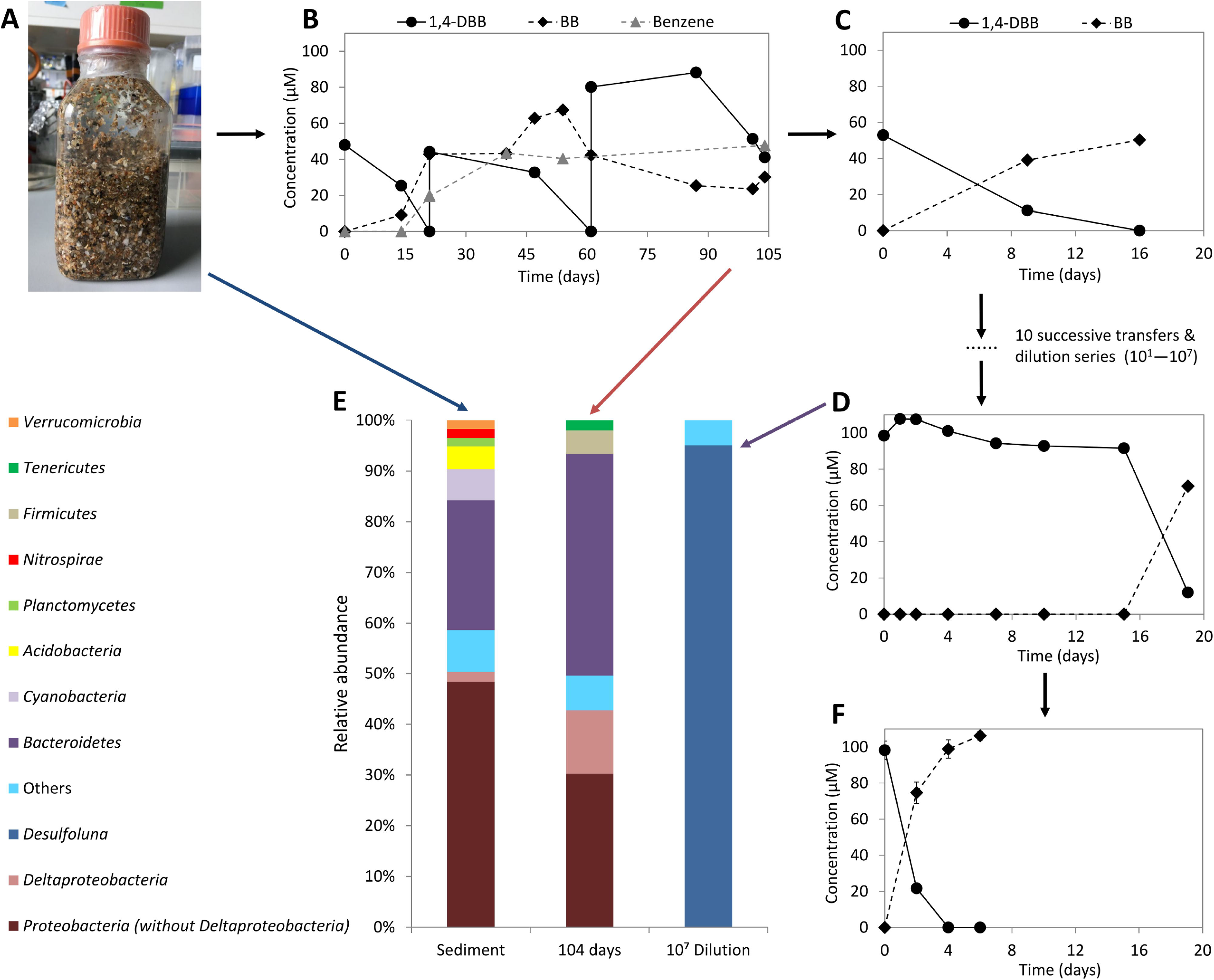
Enrichment and isolation of *D. spongiiphila* DBB. Intertidal sediment mainly composed of shore sediment used for isolation (A). Reductive debromination of 1,4-DBB by: the original microcosms containing intertidal sediment (B), the sediment-free enrichment cultures (C), the most diluted culture (10^7^) in the dilution series (D). Phylogenetic analysis of bacterial communities in the microcosms from the shore sediment at time zero (left), the original 1,4-DBB debrominating enrichment culture after 104 days incubation (middle) and the 10^7^ dilution series culture (right) (E). Reductive debromination of 1,4-DBB by the isolated pure culture (F). Sediment enrichment culture and sediment-free transfer cultures (B—D) were prepared in single bottles. Pure cultures (F) were prepared in duplicate bottles. Points and error bars represent the average and standard deviation of samples taken from the duplicate cultures. Phylogenetic data are shown at phylum level, except *Deltaproteobacteria* shown at class level and *Desulfoluna* at genus level. Taxa comprising less than 1% of the total bacterial community are categorized as Others’.

Single colonies were observed in roll tubes with 0.8% low-melting agarose after 15 days of incubation. Among the six single colonies randomly selected and transferred to liquid media, one showed 1,4-DBB debromination (Fig. 1F) which was again subjected to the roll tube isolation procedure to ensure purity. The final isolated strain was designated DBB.

### Characterization of the *Desulfoluna* strains

Cells of strain DBB were slightly curved rods with a length of 1.5 to 3 μm and a diameter of 0.5 μm as revealed by SEM (Fig. S1A, S1B), which was similar to strain AA1^⊤^ (Fig. S1C) and MSL71^⊤^ (Fig. S1D). In contrast to strain AA1^⊤^ [20], but similar to strain MSL71^⊤^ [32], strain DBB was motile when observed by light microscopy, with evident flagella being observed by SEM (Fig. S1A, B).

The cellular fatty acid profiles of the three strains consisted mainly of even-numbered saturated and mono-unsaturated fatty acids (Table S2).

Strain DBB used lactate, pyruvate, formate, malate and butyrate as electron donors for sulfate reduction (Table 1). Lactate was degraded to acetate, which accumulated without further degradation, and sulfate was reduced to sulfide (Fig. S1A). In addition, sulfite and thiosulfate were utilized as electron acceptors with lactate as the electron donor (Table 1). Sulfate and 1,4-DBB could be concurrently utilized as electron acceptors by strain DBB (Fig. S2). Independent of the presence of sulfate in the medium, strain DBB stoichiometrically debrominated 1,4-DBB to bromobenzene (BB), and 2-bromophenol (2-BP), 4-bromophenol (4-BP), 2,4-bromophenol (2,4-DBP), 2,6-DBP, 2,4,6-tribromophenol (2,4,6-TBP), 2-iodophenol (2-IP) and 4-iodophenol (4-IP) to phenol (Table 1) using lactate as the electron donor. Hydrogen was not used as an electron donor for 1,4-DBB debromination (data not shown). Strain DBB was unable to dehalogenate the tested chlorinated aromatic compounds and several other bromobenzenes listed in Table 1. This is in accordance with the dehalogenating activity reported for strain AA1^⊤^ that was unable to use chlorinated aromatic compounds as electron acceptors [20]. The majority of the known organohalogens from marine environments are brominated [1] and hence marine OHRB may be less exposed to organochlorine compounds in their natural habitats. For instance, strain AA1^⊤^ was isolated from the sponge *Aplysina aerophoba* [20] in which organobromine metabolites can account for over 10% of the sponge dry weight [53].

**Table 1.**
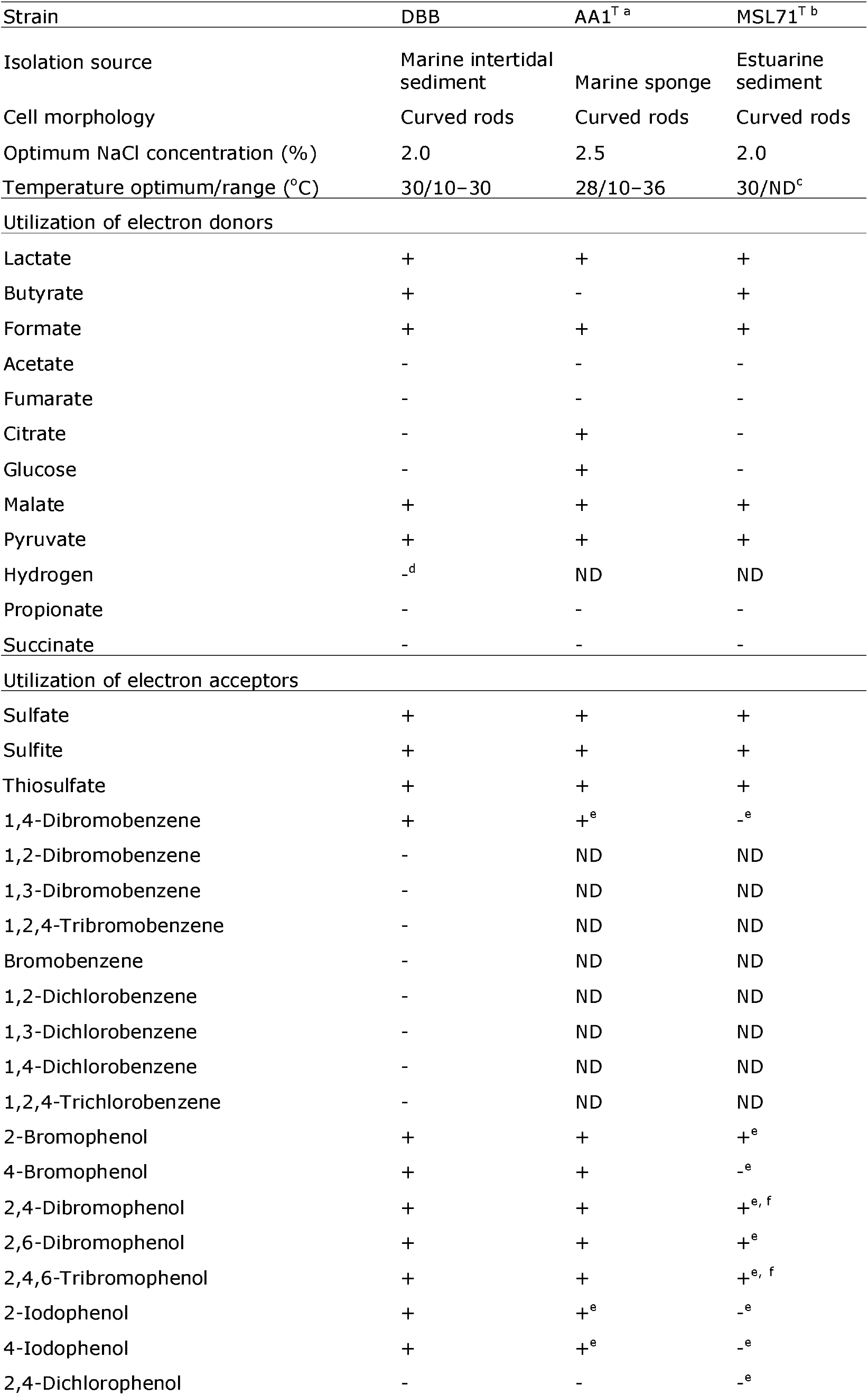
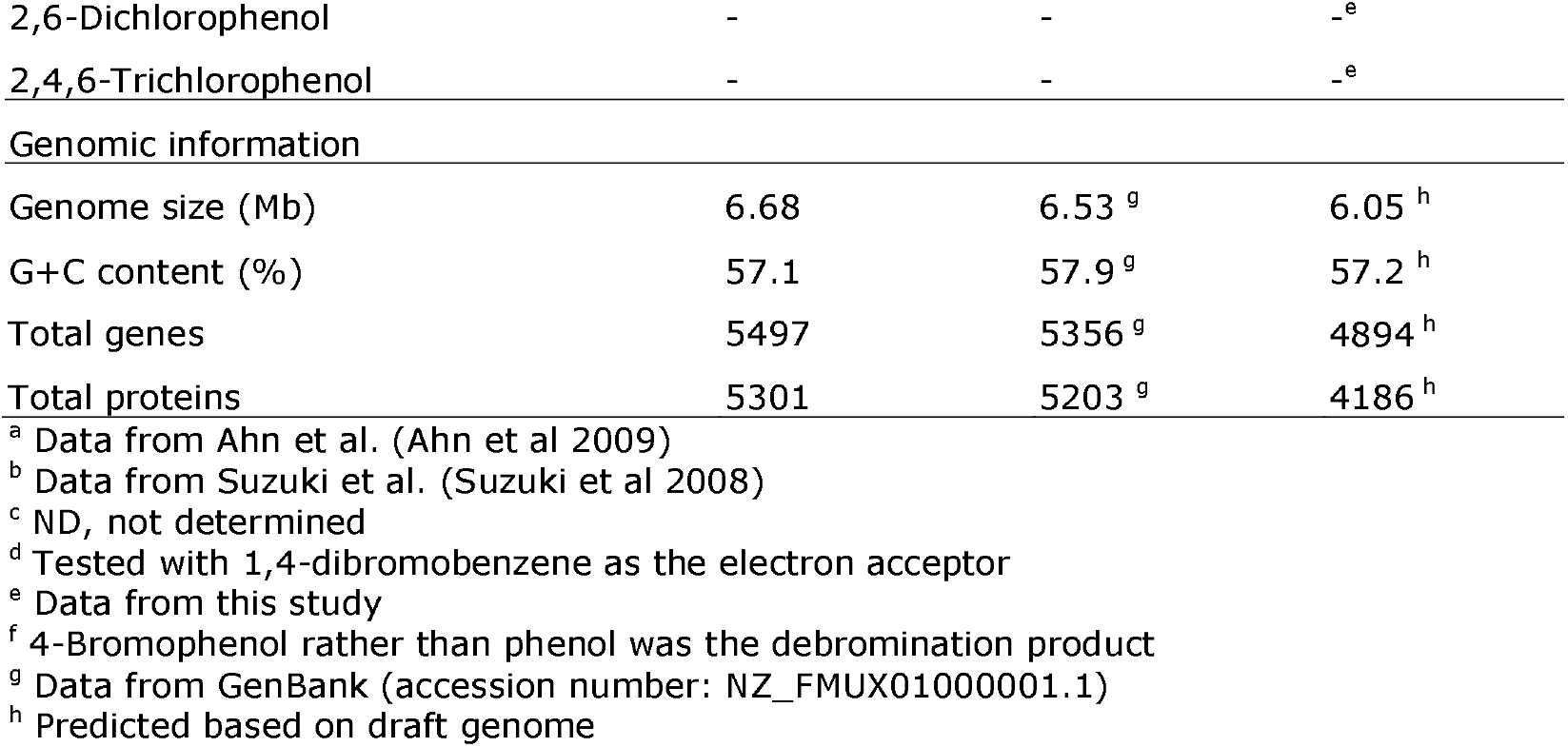
Physiological and genomic properties of *Desulfoluna* strains

| Strain | DBB | AA1<sup>T a</sup> | MSL71<sup>T b</sup> |
| --- | --- | --- | --- |
| Isolation source | Marine intertidal sediment | Marine sponge | Estuarine sediment |
| Cell morphology | Curved rods | Curved rods | Curved rods |
| Optimum NaCl concentration (%) | 2.0 | 2.5 | 2.0 |
| Temperature optimum/range (°C) | 30/10–30 | 28/10–36 | 30/ND<sup>c</sup> |
| Utilization of electron donors |  |  |  |
| Lactate | + | + | + |
| Butyrate | + | - | + |
| Formate | + | + | + |
| Acetate | - | - | - |
| Fumarate | - | - | - |
| Citrate | - | + | - |
| Glucose | - | + | - |
| Malate | + | + | + |
| Pyruvate | + | + | + |
| Hydrogen | -<sup>d</sup> | ND | ND |
| Propionate | - | - | - |
| Succinate | - | - | - |
| Utilization of electron acceptors |  |  |  |
| Sulfate | + | + | + |
| Sulfite | + | + | + |
| Thiosulfate | + | + | + |
| 1,4-Dibromobenzene | + | +<sup>e</sup> | -<sup>e</sup> |
| 1,2-Dibromobenzene | - | ND | ND |
| 1,3-Dibromobenzene | - | ND | ND |
| 1,2,4-Tribromobenzene | - | ND | ND |
| Bromobenzene | - | ND | ND |
| 1,2-Dichlorobenzene | - | ND | ND |
| 1,3-Dichlorobenzene | - | ND | ND |
| 1,4-Dichlorobenzene | - | ND | ND |
| 1,2,4-Trichlorobenzene | - | ND | ND |
| 2-Bromophenol | + | + | +<sup>e</sup> |
| 4-Bromophenol | + | + | -<sup>e</sup> |
| 2,4-Dibromophenol | + | + | +<sup>e, f</sup> |
| 2,6-Dibromophenol | + | + | +<sup>e</sup> |
| 2,4,6-Tribromophenol | + | + | +<sup>e, f</sup> |
| 2-Iodophenol | + | +<sup>e</sup> | -<sup>e</sup> |
| 4-Iodophenol | + | +<sup>e</sup> | -<sup>e</sup> |
| 2,4-Dichlorophenol | - | - | -<sup>e</sup> |
| 2,6-Dichlorophenol | - | - | -<sup>e</sup> |
| 2,4,6-Trichlorophenol | - | - | -<sup>e</sup> |
| Genomic information |  |  |  |
| Genome size (Mb) | 6.68 | 6.53<sup>g</sup> | 6.05<sup>h</sup> |
| G+C content (%) | 57.1 | 57.9<sup>g</sup> | 57.2<sup>h</sup> |
| Total genes | 5497 | 5356<sup>g</sup> | 4894<sup>h</sup> |
| Total proteins | 5301 | 5203<sup>g</sup> | 4186<sup>h</sup> |

<sup>a</sup> Data from Ahn et al. (Ahn et al 2009)
<sup>b</sup> Data from Suzuki et al. (Suzuki et al 2008)
<sup>c</sup> ND, not determined
<sup>d</sup> Tested with 1,4-dibromobenzene as the electron acceptor
<sup>e</sup> Data from this study
<sup>f</sup> 4-Bromophenol rather than phenol was the debromination product
<sup>g</sup> Data from GenBank (accession number: NZ\_FMUX01000001.1)
<sup>h</sup> Predicted based on draft genome

### Genomic and phylogenetic characterization of the *Desulfoluna* strains

The genome of strain DBB is closed and consists of a single chromosome with a size of 6.68 Mbp (Fig. S3). The genome of strain AA1^⊤^ (GenBank accession number: NZ_FMUX01000001.1) and strain MSL71T (sequenced in this study) are draft genomes with similar G+C content (Table 1). The average nucleotide identity (ANI) of the DBB genome to AA1^⊤^ and MSL71^⊤^ genomes was 98.5% and 85.9%, respectively. This indicates that DBB and AA1^⊤^ strains belong to the same species of *D. spongiiphila* [54]. 16S rRNA gene and protein domain-based phylogenetic analyses with other genera of the *Desulfobacteraceae* placed *Desulfoluna* strains in a separate branch of the corresponding phylogenetic trees (Fig. 2). Whole genome alignment of strains DBB, AA1^⊤^ and MSL71^⊤^ revealed the presence of 11 locally colinear blocks (LCBs) with several small regions of inversion and rearrangement (Fig. S4). A site-specific recombinase gene (DBB_14420) was found in one of the LCBs. The same gene was also found in the corresponding inversed and rearranged LCBs in AA1^⊤^ (AA1_11599) and MSL71^⊤^ (MSL71_ 48620), suggesting a role of the encoded recombinase in genomic rearrangement in the *Desulfoluna* strains.

**Figure 2.**
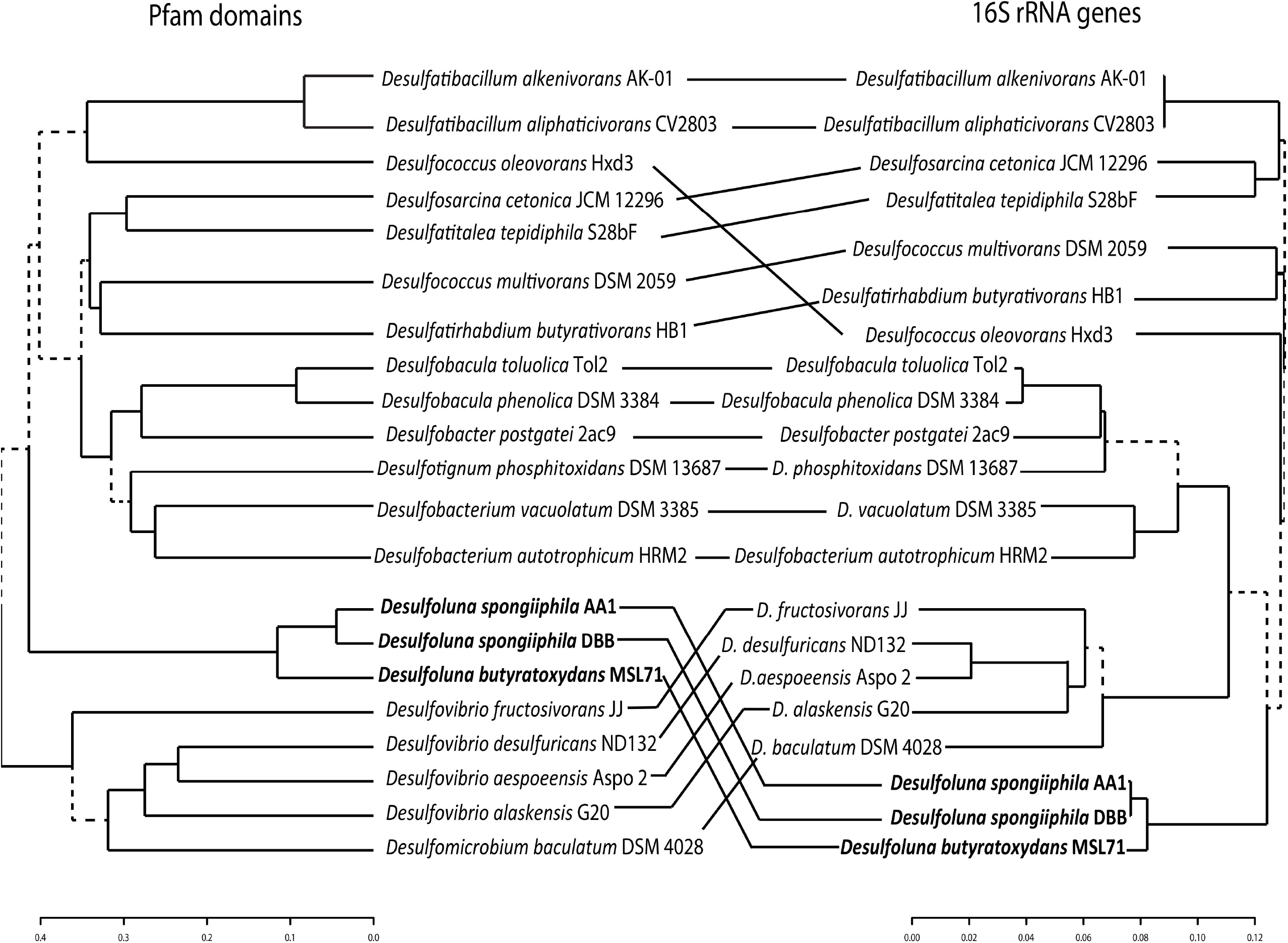
Phylogenetic tree based on 16S rRNA gene sequence and protein domain analyses. A comparison is included as horizontal lines between the two trees, showing the position of strain DBB relative to other strains belonging to the family *Desulfobacteraceae* as well as several *Desulfovibrio* strains. The “unique” nodes between the 16S rRNA gene- and domain-based tree are indicated with dashed lines. Genomes (Table S3) were selected based on the phylogenetic tree of the family *Desulfobacteraceae* [86].

### Comparison of the *rdh* gene region of the *Desulfoluna* strains

Similar to strain AA1^⊤^[21], the genomes of strains DBB and MSL71^⊤^ also harbor three *rdhA* genes. The amino acid sequences of the RdhA homologs in DBB share >99% identity to the corresponding RdhAs in AA1^⊤^, and 80—97% identity with the corresponding RdhAs in MSL71^⊤^ (Fig 3). However, the three distinct RdhA homologs in the *Desulfoluna* strains share low identity (20—30%) with each other and form three distant branches in the phylogenetic tree of RdhAs [18], and cannot be grouped with any of the currently known RdhA groups (Fig. S5). Therefore, we propose three new RdhA homolog groups, RdhA1 including DBB_38400, AA1_07176 and MSL71_22580; RdhA2 including DBB_36010, AA1_02299 and MSL71_20560; RdhA3 including DBB_45880, AA1_11632 and MSL71_30900 (Fig 3, Fig. S5).

**Figure 3.**
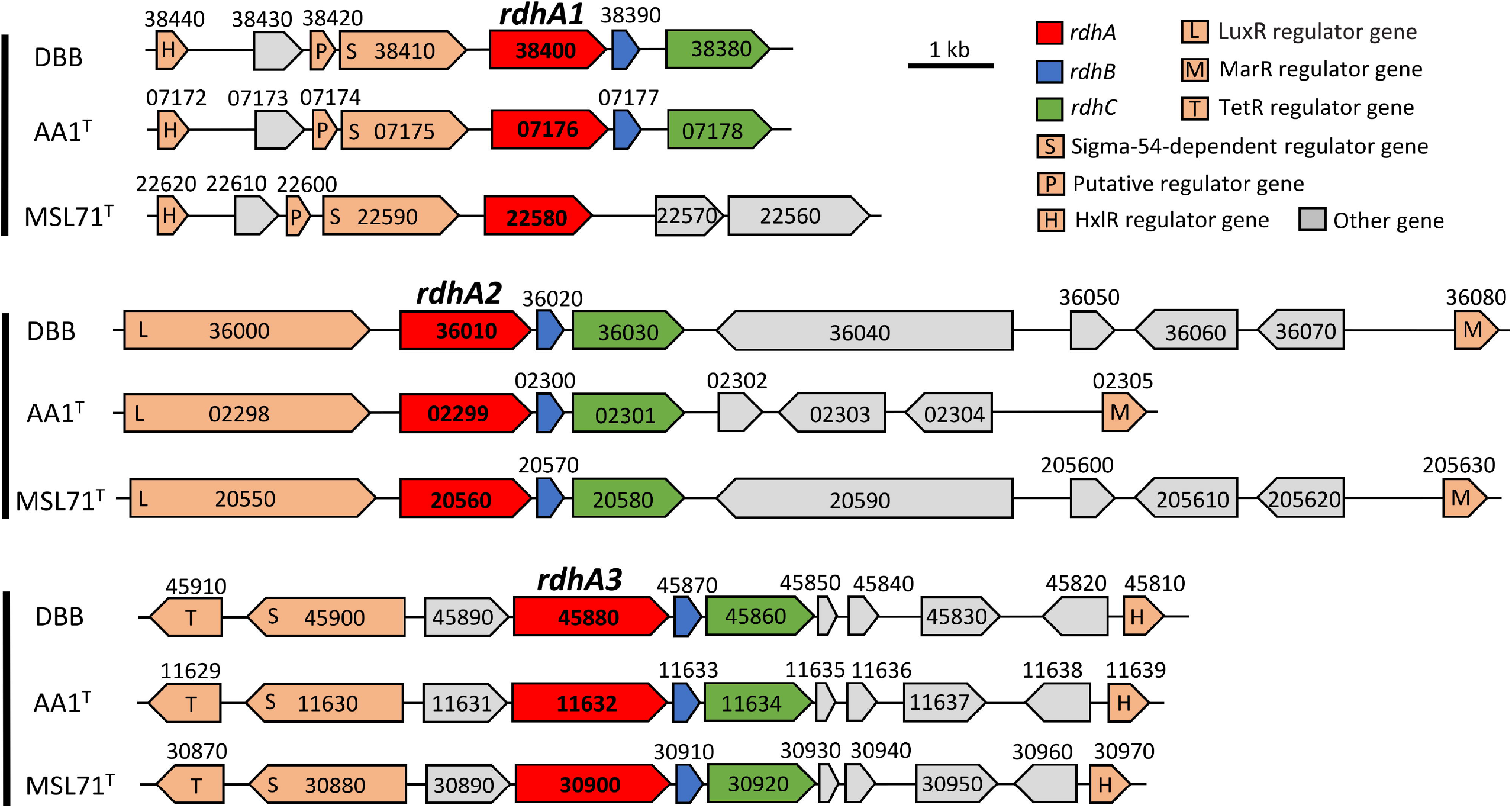
Comparison of the *rdh* gene clusters in *D. spongiiphila* DBB, *D. spongiiphila* AA1^⊤^ and *D. butyratoxydans* MSL71^⊤^. Numbers indicate the locus tags of the respective genes.

The *rdh* gene clusters in DBB and MSL71^⊤^ show a similar gene order to the corresponding *rdh* gene clusters in AA1^⊤^(Fig. 3), except that the *rdhA1* gene cluster of MSL71^⊤^ lacks *rdhB* and *rdhC*. Genes encoding sigma-54-dependent transcriptional regulators in the *rdhA1* and *rdhA3* gene clusters of AA1^⊤^ [21], were also present in the corresponding gene clusters of DBB and MSL71^⊤^ (Fig. 3). Likewise, genes encoding the LuxR and MarR-type regulators are present up- and downstream of the *rdhA2* gene clusters of DBB and MSL71^⊤^ in line with the organization of the *rdhA2* gene cluster of AA1^⊤^ (Fig. 3). This may indicate similar regulation systems of the *rdh* genes in the *Desulfoluna* strains studied here. The conserved motifs from known RDases (RR, C1–C5, FeS1, and FeS2) [55, 56] are also conserved among all the RdhAs of the *Desulfoluna* strains, except for RdhA1 of MSL71^⊤^ which lacks the RR motif (Fig. S6). This may indicate a cytoplasmic localization and a non-respiratory role of the RdhA1 in strain MSL71^⊤^ [6].

### OHR metabolism of *D. butyratoxydans* MSL71^⊤^

Guided by the genomic potential of strain MSL71^⊤^ for OHR, physiological experiments in this study indeed confirmed that strain MSL71^⊤^ is capable of using 2-BP, 2.4-DBP, 2,6-DBP and 2,4,6-TBP as electron acceptors with lactate as the electron donor. Similar to DBB and AA1^⊤^, chlorophenols such as 2,4-DCP, 2,6-DCP and 2,4,6-TCP were not dehalogenated by strain MSL71^⊤^ (Table 1). In contrast to strains DBB and AA1^⊤^, strain MSL71^⊤^ was unable to debrominate 1,4-DBB and 4-BP. Hence, debromination of 2.4-DBP and 2,4,6-TBP was incomplete with 4-BP as the final product rather than phenol (Table 1). Moreover, strain MSL71^⊤^ was unable to deiodinate 2-IP and 4-IP, again in contrast to strains DBB and AA1^⊤^ (Fig. S7, Table 1).

### Induction of *rdhA* genes during OHR by strain DBB

When strain DBB was grown with sulfate and 1,4-DBB with concomitant production of BB (Fig. 4A), its *rdhA1* gene showed significant up-regulation (60-fold) at 24 h, reached its highest level (120-fold) at 48 to 72 h, and then decreased (Fig. 4B). In contrast, no significant up-regulation of *rdhA2* or *rdhA3* was noted, suggesting that RdhA1 mediates 1,4-DBB debromination. Accordingly, RdhA1 was found in the proteome of the LSD growth condition but not in that of the LS condition (Table S4, Dataset S1 and S2). When strain DBB was grown with sulfate and 2,6-DBB, both *rdhA1* and *rdhA3* were significantly up-regulated and reached their highest level at 4 h (65- and 2000-fold, respectively, Fig. 4D). However, *rdhA3* was the dominant gene at 8 h (Fig. 4D), after which 2-BP was debrominated to phenol (Fig. 4C) indicating a role of RdhA3 in 2,6-DBP and 2-BP debromination by strain DBB. A previous transcriptional study of the *rdhA* genes in strain AA1^⊤^ during 2,6-DBP debromination also showed a similar induction of its *rdhA3* [21].

**Figure 4.**
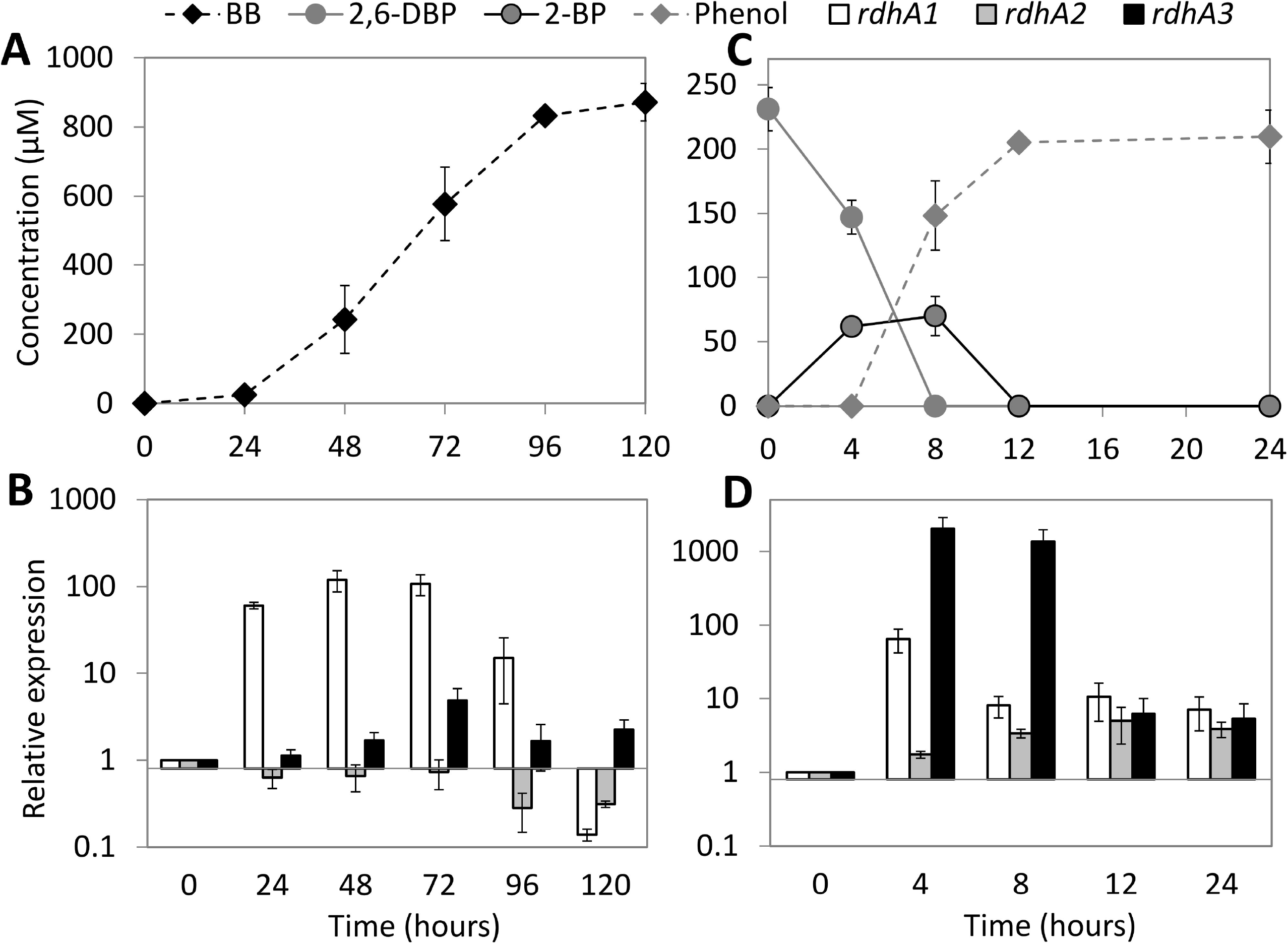
Debromination of 1,4-DBB (A) and 2,6-DBP (C) by *D. spongiiphila* DBB and relative induction of its three *rdhA* genes during debromination of 1,4-DBB (B) and 2,6-DBP (D). Error bars in panels A and C indicate the standard deviation of two random cultures analyzed out of 10 replicates. The concentration of 1,4-DBB (> 0.1 mM) could not be accurately measured due to large amount of undissolved compound and hence was not plotted. Error bars in panels B and D indicate standard deviation of triplicate RT-qPCRs performed on samples withdrawn from duplicate cultures at each time point (n = 2 × 3).

### Corrinoid biosynthesis in *Desulfoluna* strains

Most known RDases depend on corrinoid cofactors such as cyanocobalamin for dehalogenation activity [10]. Both strains DBB (this study) and AA1^⊤^ [21] were capable of OHR in the absence of externally added cobalamin. With one exception (*cbiJ*), the genomes of the *Desulfoluna* strains studied here harbor all genes necessary for *de novo* anaerobic corrinoid biosynthesis starting from glutamate (Table S5). The genes for cobalamin biosynthesis from precorrin-2 are arranged in one cluster (DBB_3730—3920, AA1_12810—12829, MSL71_49290—49480) including an ABC transporter (*btuCDF)* for cobalamin import (Fig. 5). Another small cobalamin-related gene cluster was detected in the *Desulfoluna* genomes (DBB_52170–52260, AA1_10815–10826, MSL71_44540–44630), which includes genes coding for the outer membrane corrinoid receptor BtuB and a second copy of the corrinoid-transporter BtuCDF plus another BtuF. Additionally, cobaltochelatase CbiK as well as a putative cobaltochelatase CobN are encoded in this gene cluster. The latter is usually involved only in the aerobic cobalamin biosynthesis pathway, and its function in *Desulfoluna* strains is unknown. Three of the proteins encoded by DBB_3730—3920 (Cbik: 3730, CbiL: 3790, CbiH: 3850) were detected in the proteome of cells grown under both the LS and LSD conditions (Table S4, Dataset S1). The abundance of the cobalamin biosynthesis proteins was not significantly different between LS and LSD conditions (Table S4, Dataset S1 and S2), except for the tetrapyrrole methylase CbiH encoded by DBB_3850 that was significantly more abundant in LSD cells (Table S4, Dataset S1). The detection of cobalamin biosynthesis proteins in the absence of 1,4-DBB in LS condition could be due to the synthesis of corrinoid-dependent enzymes in the absence of an organohalogen. Accordingly, three corrinoid-dependent methyltransferase genes (encoded by DBB_7090, 43520, 16050) were detected in the proteomes, which might be involved in methionine, methylamine or o-demethylation metabolism. This might also indicate a constitutive expression of the corresponding genes, in contrast to the organohalide-induced cobalamin biosynthesis in *Sulfurospirillum multivorans* [57].

**Figure 5.**
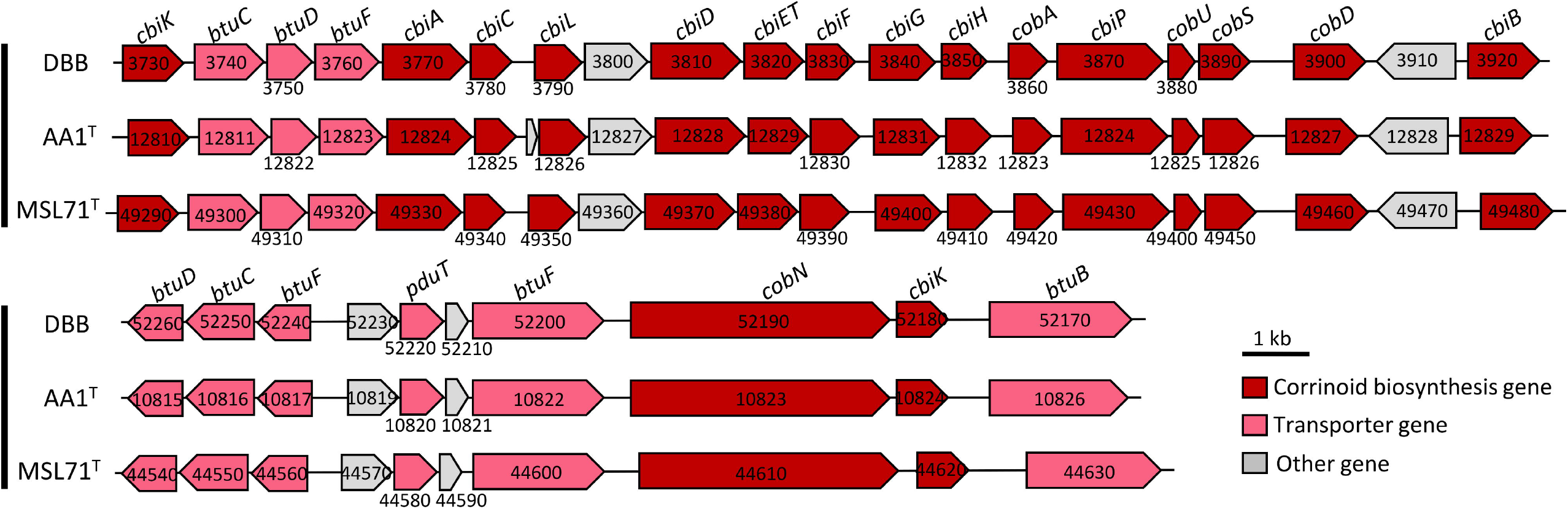
Corrinoid biosynthesis and transporter gene clusters of *Desulfoluna* strains. Numbers indicate the locus tags of the respective genes. The corresponding enzymes encoded by the genes and their functions in corrinoid biosynthesis are indicated in Table S4.

### Sulfur metabolism and impact of sulfate and sulfide on debromination by *Desulfoluna* strains

All three strains were capable of using sulfate, sulfite, and thiosulfate as the terminal electron acceptors (Table 1). Four sulfate permease genes are present in the genomes of the *Desulfoluna* strains (Table S6), and one of the sulfate permeases (DBB_22290) was detected in DBB cells grown under LS and LSD conditions (Table S4, Dataset S2). The genes involved in sulfate reduction, including those encoding sulfate adenylyltransferase (Sat), APS reductase (AprBA) and dissimilatory sulfite reductase (DsrAB), were identified in the genomes of all three strains (Table S6). The corresponding proteins were detected in DBB cells grown under both LS and LDS conditions (Fig. 6, Table S4) with AprBA, disulfite reductase (DsrMKJOP) and Sat among the most abundant proteins in both, soluble and membrane fractions (Dataset S1 and S2). Tetrathionate reductase encoding genes (*ttrA*) were found only in the genomes of strains DBB and AA1^⊤^. Interestingly, thiosulfate reductase genes were not found in any of the three genomes, whereas all strains can use thiosulfate as the electron acceptor (Table 1). *Desulfitobacterium metallireducens* was also reported to reduce thiosulfate despite lacking a known thiosulfate reductase gene [58, 59], suggesting the existence of a not-yet-identified gene encoding a thiosulfate reductase [59]. Possible alternatives are genes encoding rhodanese-like protein (RdlA) (Table S6) [60] or the three-subunit, periplasmic molybdopterin oxidoreductase (Table S6), as a putative polysulfide reductase (Psr) [61].

**Figure 6.**
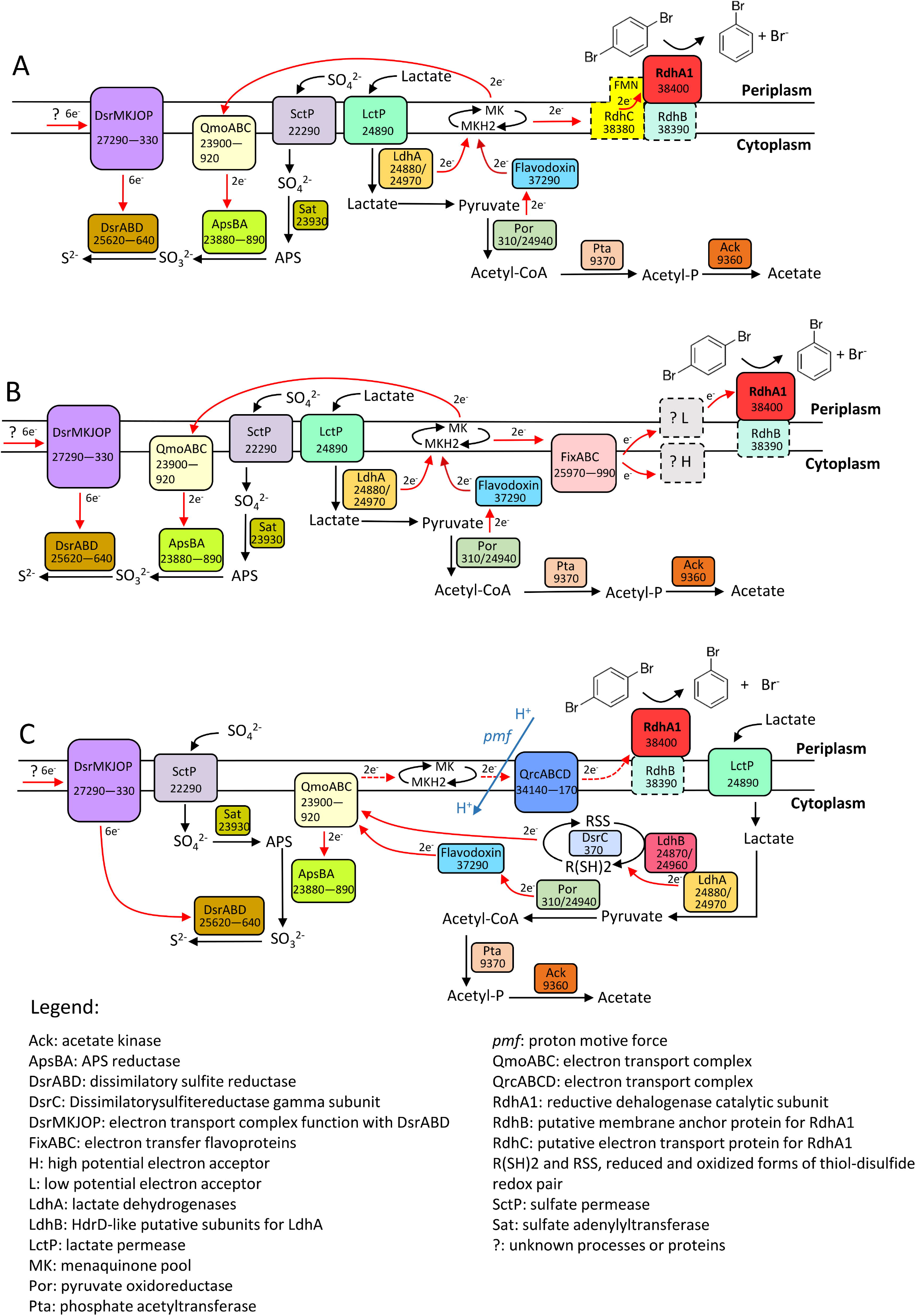
Proposed electron transport pathways with OHR mediated by RdhC (A), Fix complex (B), Qmo/Qrc complexes (C) in *D. spongiiphila* DBB grown on lactate and sulfate (LS) and lactate, sulfate and 1,4-DBB (LSD). Corresponding gene locus tags are given for each protein. Log protein abundance ratios between LSD and LS grown cells are indicated next to the gene locus tag. Proteins shown in dashed line square were not detected under the tested conditions. Probable electron flow path is shown in red arrows, and the dashed red arrows indicate reverse electron transport.

Sulfate and sulfide are known inhibitors of many OHRB [30, 62, 63]. However, debromination of 2,6-DBP was not affected in *Desulfoluna* strains in the presence of up to 20 mM sulfate (Fig. S8B, D, F), and sulfate and 2,6-DBP were reduced concurrently (Fig. S8). This is similar to some other *Deltaproteobacteria* [23], but in contrast to *D. tiedjei* which preferentially performs sulfate reduction over OHR with concomitant down-regulation of *rdh* gene expression [30]. Moreover, sulfide, an RDase inhibitor in *D. tiedjei* [64] and *Dehalococcoides mccartyi* strains [28, 29], did not impact 2,6-DBP debromination by *Desulfoluna* strains at a concentration of 10 mM (Fig. S9A—F). However, debromination was delayed in the presence of 20 mM sulfide, and no debromination was noted in the presence of 30 mM sulfide (Fig. S9G—L). This high resistance to sulfide was not reported before for the known OHRB, and is also rare among sulfate-reducing bacteria [65], and may confer an ecological advantage to these sulfate-reducing OHRB. Although hydrogen sulfide can be oxidized abiotically or serve as electron donor for sulfide-oxidizing microorganisms [66], naturally sustained and high concentrations of hydrogen sulfide are found in some marine environments [67].

### Electron transport chains of strain DBB

Two lactate dehydrogenases (LdhA-1/2, DBB_24880/24970) with HdrD-like putative iron-sulfur subunits (LdhB-1/2, DBB_24870/24960) were found in the proteome of DBB cells grown under LS and LSD conditions. Similar Ldhs were reported to be essential for the growth of *Desulfovibrio alaskensis* G20 with lactate and sulfate [68]. Similar to *D. alaskensis* G20 and *D. vulgaris* strain Hildenborough [68, 69], the two Ldhs were encoded by an organic acid oxidation gene cluster (DBB_24870—24970) including genes encoding lactate permease (DBB_24890), the Ldhs and pyruvate oxidoreductase (Por, DBB_24940). Based on previous studies with *D. vulgaris* Hildenborough [70], the electron transport pathway in strain DBB with lactate and sulfate could take one of the following routes: the Ldh’s either reduce menaquinone directly [70], or transfer electrons via the HdrD-like subunit [71] and DsrC (DBB_370, a high redox potential electron carrier with disulfide/dithiol (RSS/R(SH)2)) to QmoA [72]. The pyruvate produced by lactate oxidation is further oxidized by Por (DBB_310/24940), and the released electrons are carried/transferred by a flavodoxin (DBB_37290). From there, the electrons from the low-potential ferredoxin and the electrons from the high-potential (disulfide bond) DsrC could be confurcated to QmoABC, which reduces menaquinone (Fig. 6A, B). The electrons are then transferred from menaquinol to the APS reductase (AprBA, DBB_23880—890) which is, together with three other enzyme complexes (Sat, encoded by DBB_23930, DsrABD, DBB_25620—640, and DsrMKJOP, DBB_27290—330), responsible for the sulfate reduction cascade (Santos et al., 2015).

The electron transport chain from Ldh to menaquinones or QmoABC during OHR is likely shared with sulfate reduction. Electron transport from menaquinol (*E*^0’^ = −75 mV) to the RDase (*E*^0’^ (CoII/CoI) ≈ −360 mV) is thermodynamically unfavorable (Schubert et al., 2018), and the proteins involved to overcome this barrier have not been identified and most likely are not the same in different organohalide-respiring bacterial genera. Based on the genomic and proteomic analyses of strain DBB, we identified several possible electron transfer proteins connecting the menaquinone pool and RdhA1. The first is the membrane-integral protein RdhC1 (encoded by DBB_38380, Fig. 3), a homolog of proteins previously proposed to function as transcriptional regulator for *rdhAB* gene expression in *Desulfitobacterium dehalogenans* [73]. However, a recent study on PceC from *Dehalobacter restrictus* proposed a possible role for RdhC in electron transfer from menaquinones to PceA via its exocytoplasmically-facing flavin mononucleotide (FMN) cofactor [16]. RdhC in *Desulfoluna* strains also showed the conserved FMN binding motif (in particular the fully conserved threonine residue) and two CX_3_CP motifs predicted to have a role in electron transfer [16] (Fig. S10). Moreover, the five transmembrane helices of RdhC in DBB were also conserved (Fig. S11), indicating a possible function of RdhC1 in electron transfer from menaquinones to RdhA1 (Fig. 6A). However, RdhC1 was not found in our proteomic analysis, probably due to tight interaction with the membrane.

A second link between menaquinol/QmoABC and RdhA1 could be the Fix complex homolog, an electron transfer flavoprotein complex found in nitrogen-fixing microorganisms such as *Azotobacter vinelandii* and *Rhodospirillum rubrum* [74, 75]. The Fix complex is capable of using electron bifurcation to generate low-potential reducing equivalents for nitrogenase [74]. Strain DBB does not encode the minimum genes necessary for nitrogen fixation [76]. Hence, the Fix complex in DBB cells is likely linked to other cellular processes. Induction of the *fix* genes under OHR conditions was reported in other OHRB such as *Desulfitobacterium hafniense* TCE1 [77], and the corresponding Fix complex was suggested to provide low-redox-potential electrons for OHR. However, the obligate organohalide-respiring *Dehalobacter* spp., which are phylogenetically related to *Desulfitobacterium* spp., do not encode FixABC, questioning a general role of Fix complex in OHR (Türkowsky et al., 2018). In strain DBB, the abundance of FixABC (encoded by DBB_25970—990) was not higher in the cells grown under LDS as opposed to LS condition, but FixAB were among the most abundant 10% proteins in the soluble fraction (Dataset S1), indicating a potential role in electron transfer in both sulfate reduction and OHR. In this scenario, FixABC accepts two electrons from menaquinol, subsequently bifurcating them to unidentified high- and low-potential electron acceptors (Fig. 6B). The low-potential electron acceptor may also serve as an electron carrier that transfers electrons from cytoplasm-facing FixABC to the exoplasm-facing RdhA1 via an as-yet-unidentified electron carrier across the membrane [78] (Fig. 6B).

A third scenario is the involvement of QmoABC- and QrcABCD-mediated reverse electron transport (Fig. 6C), similar to the electron transport system of *D. alaskensis* G20 cultivated in syntrophic interaction with *Methanococcus maripaludis* [68]. The electron transport from menaquinol to the periplasmic hydrogenase or formate dehydrogenase in strain G20 also needs to overcome an energy barrier similar to that of OHR (redox potential of H_2_/H^+^ and formate/CO_2_ are −414 mV and −432 mV, respectively) [68]. In this scenario, lactate is oxidized to pyruvate as described above, transferring electrons to a thiol-disulfide redox pair. Pyruvate is oxidized by Por and the electrons are accepted by the flavodoxin. QmoABC then confurcates electrons from the low-potential ferredoxin and the high-potential thiol-disulfide redox pair to drive reduction of menaquinones. Electrons are transferred from menaquinol to RdhA1 via QrcABCD by reverse electron transport (Fig. 6C). The energy required for reverse electron transport is likely derived from the proton motive force mediated by QrcABCD [79]. In this scenario, QmoABC plays a key role in the metabolism of strain DBB as a link between sulfate reduction and OHR. This electron transport pathway provides a possible explanation for the increased 1,4-DBB debromination rate by DBB when sulfate is concurrently present (Fig. 1E, Fig. S1B). Hence, sulfate reduction may stimulate the electron confurcation process that is also used for OHR. Moreover, sulfate reduction can generate the proton motive force required for the reverse electron transport from QmoABC to RdhA1. Qmo and Qrc complexes are frequently found in sulfate-reducing *Deltaproteobacteria* and were proposed to be involved in energy conservation [71, 80, 81]. However, biochemical studies with sulfate-reducing OHRB are necessary to further corroborate such a reverse electron flow and the intricate relationship of electron transfer in sulfate reduction and OHR.

### Potential oxygen defense in *Desulfoluna* strains

Sulfate reducers, which have been assumed to be strictly anaerobic bacteria, not only survive oxygen exposure but also can utilize it as an electron acceptor [82, 83]. However, the response of organohalide-respiring sulfate reducers to oxygen exposure is not known. Most of the described OHRB are strict anaerobes isolated from anoxic and usually organic matter-rich subsurface environments [17]. In contrast, strain DBB was isolated from marine intertidal sediment mainly composed of shore sand (Fig. 1A), where regular exposure to oxic seawater or air can be envisaged. The genomes of the *Desulfoluna* strains studied here harbor genes encoding enzymes for oxygen reduction and reactive oxygen species (ROS) detoxification (Table S7). Particularly, the presence of a cytochrome *c* oxidase is intriguing and may indicate the potential for oxygen respiration. Accordingly, in the presence of 2% oxygen in the headspace of DBB cultures, the redox indicator resazurin in the medium turned from pink to colorless within two hours, indicating consumption/reduction of oxygen by strain DBB. Growth of strain DBB on lactate and sulfate was retarded in the presence of 2% oxygen (Fig. S12C). However, in both the presence (Fig. S12C) and absence of sulfate (Fig. S12D), slower but complete debromination of 2,6-DBP to phenol was achieved with 2% oxygen in the headspace. Neither growth nor 2,6-DBP debromination was observed with an initial oxygen concentration of 5% in the headspace (Fig. S12E, F). Such resistance of marine OHRB to oxygen may enable them to occupy niches close to halogenating organisms/enzymes that nearly all use oxygen or peroxides as reactants [84]. For instance, the marine sponge *A. aerophoba* from which *D. spongiiphila* AA1^⊤^ was isolated [20] harbors bacteria with a variety of FADH_2_-dependent halogenases [85], and produces a variety of brominated secondary metabolites [53]. Testing survival and OHR of *Desulfoluna* strains under continuous oxygen exposure and studying the mechanisms of oxygen defense as studied in *Sulfurospirillum multivorans* [11] are necessary to further unravel oxygen resistance/metabolism mechanisms in *Desulfoluna* strains.

## Conclusions

Widespread environmental contamination with organohalogen compounds and their harmful impacts to human and environmental health has been the driver of chasing OHRB since the 1970s. In addition, the environment itself is an ample and ancient source of natural organohalogens, and accumulating evidence shows widespread occurrence of *rdhA* in marine environments [6]. The previous isolation and description of strain AA1^⊤^ from a marine sponge, the isolation of strain DBB from intertidal sediment samples, and verification of the OHR potential of strain MSL71^⊤^ in this study indicate niche specialization of the members of the genus *Desulfoluna* as chemoorganotrophic facultative OHRB in marine environments rich in sulfate and organohalogens. As such, *de novo* corrinoid biosynthesis, resistance to sulfate, sulfide and oxygen, versatility in using electron donors, and the capacity for concurrent sulfate and organohalogen respiration confer an advantage to *Desulfoluna* strains in marine environments. Interestingly, approximately 10% of the sequenced deltaproteobacterial genomes, that have mostly been obtained from marine environments, contain one or multiple *rdh* genes [22, 23], and OHR metabolism was experimentally verified in three strains not previously known as OHRB [23]. These findings reinforce an important ecological role of sulfate-reducing organohalide-respiring *Deltaproteobacteria* in sulfur, halogen and carbon cycling in a range of marine environments.

## Acknowledgements

We would like to thank Johanna Gutleben and Maryam Chaib de Mares for sediment sampling, W. Irene C. Rijpstra for fatty acid analysis, and Andreas Marquardt (Proteomics Centre of the University of Konstanz) for proteomic analyses. We acknowledge the China Scholarship Council (CSC) for the support to PP and YL. The authors thank BE-BASIC funds (grants F07.001.05 and F08.004.01) from the Dutch Ministry of Economic Affairs, ERC grant (project 323009), the Gravitation grant (project 024.002.002) of the Netherlands Ministry of Education, Culture and Science and the Netherlands Science Foundation (NWO), and National Natural Science Foundation of China (project No.51709100) for funding.

