## Supplementary Information for "Organohalide-respiring *Desulfoluna* species isolated from marine environments"

**Enrichment, isolation and cultivation of strain DBB**

The sediment sampling bottles were filled with seawater to leave no headspace. For preparation of microcosms, sediment (5 g) was transferred into 120 ml bottles containing 50 ml of anoxic medium [1] and N_2_/CO_2_ (80 : 20%, 140 kPa) as the headspace. Vitamins and trace elements were added as described previously [2] except that cyanocobalamin was omitted. Lactate (5 mM) and 1,4-dibromobenzene (1,4-DBB, 50 μM) were used as the electron donor and acceptor, respectively. 1,4-DBB was added from a 10 mM stock solution dissolved in acetone. The bottles were sealed with viton stoppers and aluminium crimp caps and incubated statically in the dark at 25°C. After debromination of three spikes of 1,4-DBB, sediment-free cultures were obtained by transferring the suspensions of the enrichment culture (10% v/v) to fresh medium using the same growth condition as described above. After ten successive transfers of the sediment-free cultures, a dilution series from 10^1^ to 10^7^-fold was performed. The most diluted culture showing 1,4-DBB debromination (10^7^-fold) was then serially diluted from 10^1^- to 10^3^-fold in 25 ml roll tubes containing 10 ml medium and 0.8% low-melting point agarose (Sigma-Aldrich) and incubated in the dark at 25°C. Individual colonies were randomly picked and transferred into liquid medium to check for 1,4-DBB debromination. A culture showing debromination activity was re-isolated in roll tubes as described above to ensure the purity.

The optimum NaCl concentration for growth of strain DBB was determined in the range from 10 to 30 g/L. Using the optimal NaCl concentration (20 g/L), the following halogenated aromatic compounds were tested as electron acceptors for strain DBB with lactate (5 mM) as the electron donor and carbon source: 1,2-dibromobenzene (1,2-DBB), 1,3-dibromobenzene (1,3-DBB), 1,2,4-tribromobenzene (1,2,4-TBB), 2-bromophenol (2-BP), 4-bromophenol (4-BP), 2,4-dibromophenol (2,4-DBP), 2,6-dibromophenol (2,6-DBP), 2,4,6-tribromophenol (2,4,6-TBP), 2-iodophenol (2-IP), 4-iodophenol (4-IP), 1,2-dichlorobenzene (1,2-DCB), 1,3-dichlorobenzene (1,3-DCB), 1,4-dichlorobenzene (1,4-DCB), 1,2,4-trichlorobenzene (1,2,4-TCB), 2,4-dichlorophenol (2,4-DCP), 2,6-dichlorophenol (2,6-DCP) and 2,4,6-trichlorophenol (2,4,6-TCP). Brominated and chlorinated benzenes, 2,4,6-TBP and 2,4,6-TCP were added from 10 mM stock solutions dissolved in acetone to nominal concentrations of 100 μM in the medium. The remaining di- and mono-brominated phenols were added from 10 mM stock solutions in 0.1 N NaOH to nominal concentrations of 50—100 μM. Sulfate, sulfite and thiosulfate (5 mM) were tested as electron acceptors with 10 mM lactate as the electron donor. To test the utilization of electron donors, acetate, propionate, fumarate, malate, butyrate, lactate, pyruvate, succinate, glucose and citrate were added separately at 10 mM to the medium containing 10 mM sulfate. Utilization of hydrogen (5 mM) and formate (5 mM) as the electron donors for debromination of 1,4-DBB (100 µM) was tested in presence of acetate (5 mM) as the carbon source. To study the effect of sulfate and sulfide on debromination, sulfate (10—20 mM) or sulfide (1−30 mM) together with lactate (20−40 mM) were added to the medium containing 100 μM of 1,4-DBB or 2,6-DBP. To test the impact of oxygen on debromination, strain DBB was grown in medium without Na_2_S as the reducing agent, in presence or absence of sulfate (10 mM). The medium contained 20 mM lactate, 100 µM 2,6-DBP and 0%, 2% or 5% oxygen in the headspace.

**Cellular fatty acids analysis**

The cultures were harvested at the early stationary growth phase by centrifugation at 4,700 × g for 15 min at 4°C. Cellular fatty acids were analysed by acid hydrolysis of total cell material following a method previously described [3]. The fatty acids were identified by analysis with gas chromatography-mass spectrometry before and after derivatisation of double bonds with dimethyl disulphide to enable localization of the double bond position [3].

**qPCR assays**

Primers for amplification of the three *rdhA* genes in strain DBB were designed using the NCBI online primer design tool (<http://www.ncbi.nlm.nih.gov/tools/primer-blast/>) (Table S1). In order to prepare standards for the qPCR assays, the *rdhA* genes were PCR amplified using the following program: 95°C for 5 min, followed by 30 cycles of 95°C for 30 s, 55°C for 30 s and 72°C for 30 s, followed by a final extension at 72°C for 10 min. The *rdhA* genes were then cloned into pGEM®-T Easy Vector (Promega, WI, USA) and introduced into *E. coli* JM109 competent cells (Promega, WI, USA). Plasmid purification and preparation of the dilution series of the qPCR standards (from 10^1^ to 10^8^ copies/µl) were done as described earlier [4]. qPCRs were performed using the iQ SYBR Green supermix (Bio-Rad, CA, USA). The qPCR program was: 95°C for 10 min, followed by 40 cycles of 95°C for 15 s, 60°C for 30 s and 72°C for 30 s. Melting curves were measured from 65°C to 95°C with increments of 0.5°C and 10 s at each step. Transcription of the *rdhA* genes was determined using cDNA as the template. The transcript levels were calculated by relative quantification using the 2^-ΔΔCq^ method with the 16S rRNA gene as the reference gene [5, 6]. Gene expression data was normalized to values observed at the 0 h time point, at which 1,4-DBB or 2,6-DBP were initially amended [5]. A relative expression difference higher than 10-fold was arbitrarily set as representing significant induction [7].

**Protein extraction and proteomic analysis**

Protein was extracted from 100 ml culture of strain DBB grown with lactate (20 mM)/sulfate (10 mM) and lactate (20 mM)/sulfate (10 mM)/1,4-DBB (100 µM); triplicate samples were prepared for each condition. Cells were collected by centrifugation at 4500 × g for 20 min at 4°C. The cells were then re-suspended in 1 ml 100 mM Tris-HCl buffer (pH 7.5) containing 10 µl protease inhibitor (Halt Protease Inhibitor Cocktail; Thermo Fisher Scientific, Rockford, USA). Cells were lysed by sonication using a Branson sonifier (Branson, CT, USA) equipped with a 3 mm tip by six pulses of 30 s with 30 s rest in between of each pulse. Cell debris was removed by centrifugation at 10,000 g for 10 min at 4°C. The protein concentration of the cell-free extracts (CFE) was determined using the Bradford assay [8]. The total-proteomics samples were prepared and the analyses were done as described by Burrichter et al. [9]. Total protein (200 µg) in CFE was purified through SDS-PAGE until the proteins had entered the stacking gel (without any separation); the Coomassie-stained total-protein bands were excised and then subjected to peptide fingerprinting-mass spectrometry (see below). For analysis of proteins associated to the membrane, the membrane fragments in the CFE were separated by ultracentrifugation at 104,000 g for 35 min at 4°C; the membrane pellet was solubilized in SDS-PAGE loading dye and the proteins were also purified by SDS-PAGE and the Coomassie-stained total-protein bands were excised, as described above. The total-protein bands excised from SDS-PAGE gels were subjected to peptide fingerprinting-mass spectrometry at the Proteomics Facility of the University of Konstanz ([www.proteomics-facility.uni-konstanz.de](http://www.proteomics-facility.uni-konstanz.de)) [9]. Each sample was analyzed twice on a Orbitrap Fusion with EASY-nLC 1200 (Thermo Fisher Scientific) and tandem mass spectra were searched against an appropriate protein database of strain DBB using Mascot (Matrix Science) and Proteome Discoverer V1.3 (Thermo Fisher Scientific) with “Trypsin” enzyme cleavage, static cysteine alkylation by chloroacetamide, and variable methionine oxidation [9].

**Analytical methods**

The column temperature program of the GC-FID was: 40°C hold for 2 min, followed by an increase of 6°C min^−1^ to 100°C and hold for 2 min, followed by further increase at 10°C min^−1^ to 225°C and hold for 2 min. The program for benzene measurement was as described earlier [10]. The wavelength of the UV detector of the HPLCs was 210 nm. The mobile phases for the Thermo Scientific Accela HPLC System were 0.1% formic acid in water (eluent A) and 0.1% formic acid in acetonitrile (eluent B). The mobile phase for the ThermoFisher Scientific SpectraSYSTEM™ HPLC was 0.01 N H_2_SO_4_. Halogenated phenols and phenol were analyzed using a three-step gradient profile consisting of: i) 90% eluent A and 10% eluent B for 2 min, ii) 90−20% eluent A and 10−80% eluent B for 14 min and hold at 20% eluent A and 80% eluent B for 3 min, iii) followed by 20—90% eluent A and 80—10% eluent B for 1 min. The ions were analyzed using a three-step gradient profile consisting of 1 mM KOH for 1 min, 1—40 mM KOH for 14 min and hold at 40 mM KOH for 4 min, followed by 40−1 mM KOH for 4.5 min.

**Supplementary Figures**

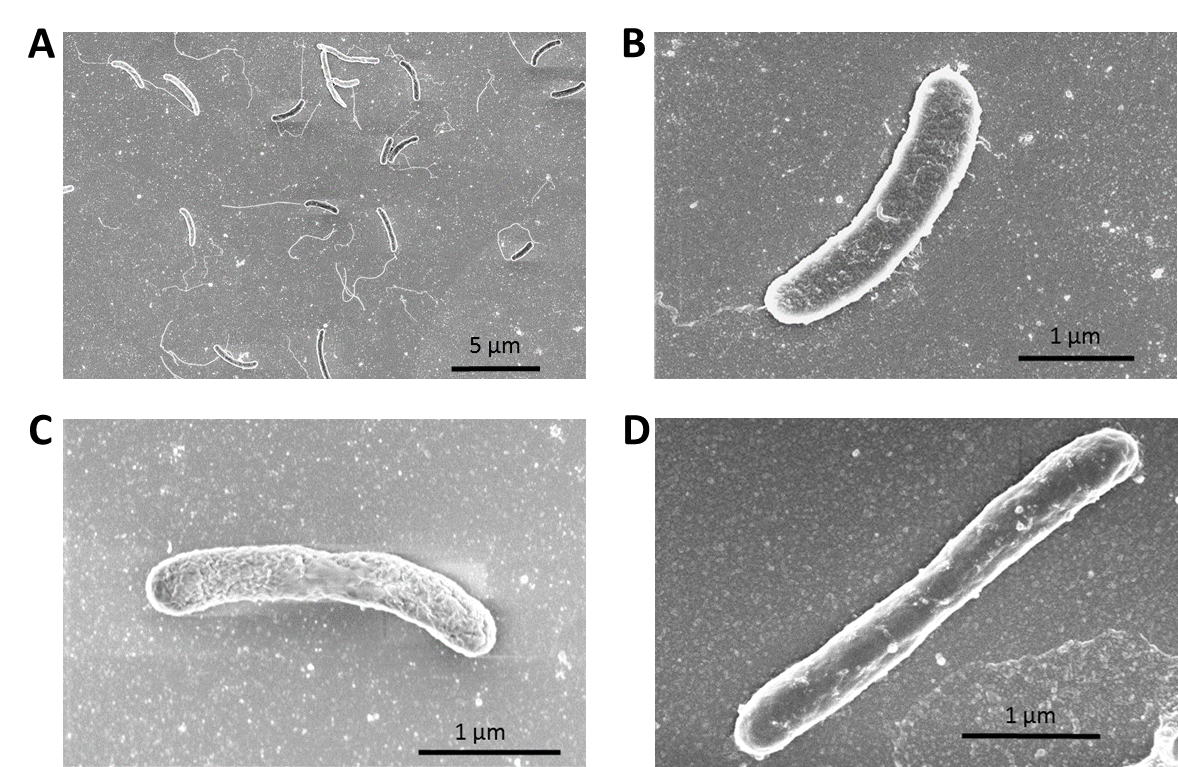

**Fig. S1** Scanning electron micrograph of *D. spongiiphila* DBB (A and B), *D. spongiiphila* AA1^T^ (C) and *D. butyratoxydans* MSL71^T^ (D).

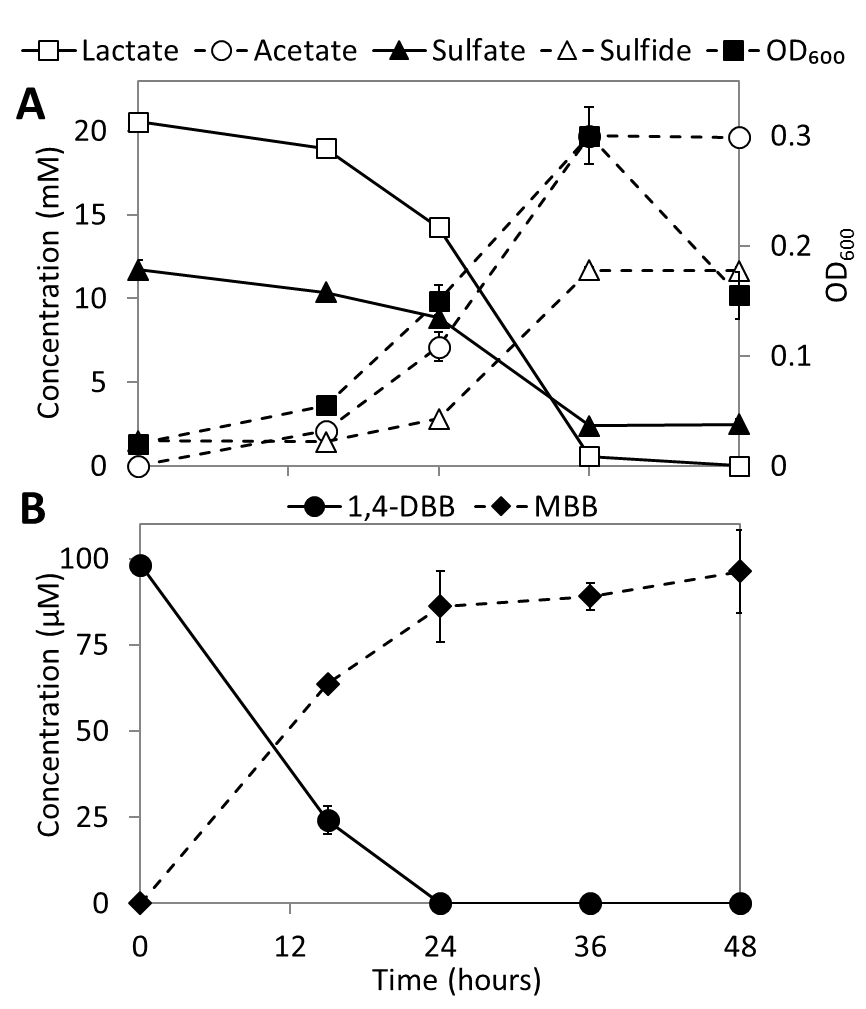

**Fig. S2** Concurrent 1,4-DBB debromination and sulfate reduction by strain DBB with lactate as the electron donor. Points and error bars represent the average and standard deviation of samples taken from duplicate cultures.

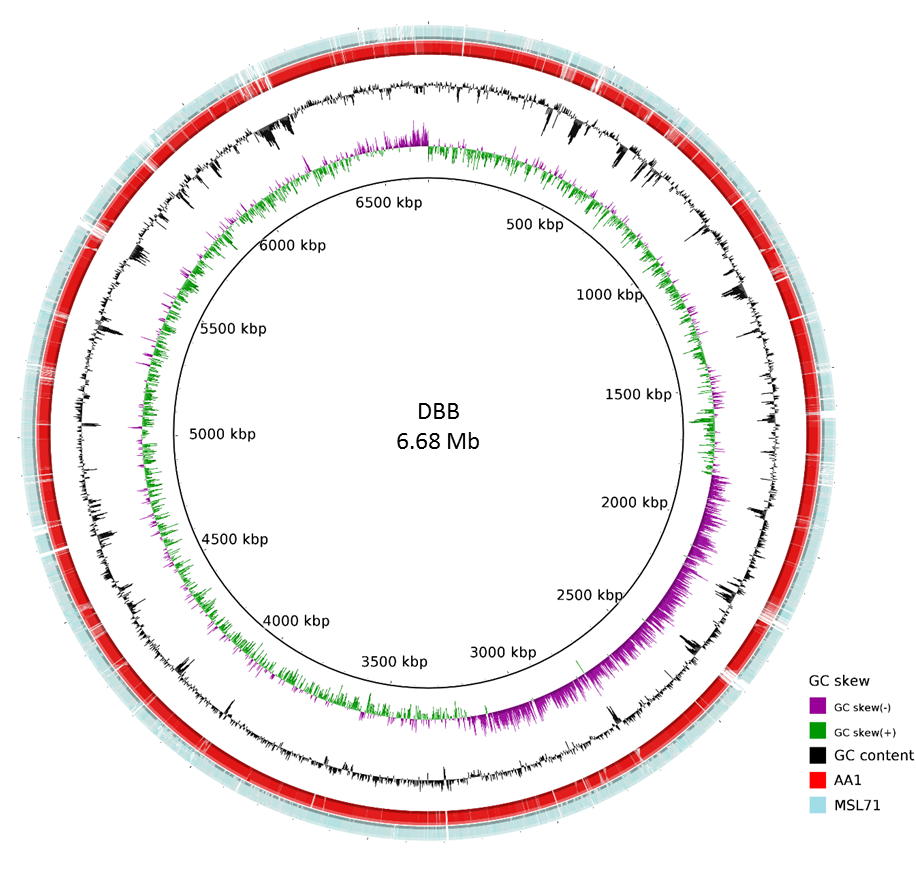

**Fig. S3** Circular representation of the genome sequence of *D. spongiiphila* DBB in comparison with the genomes of *D. spongiiphila* AA1^T^ and *D. butyratoxydans* MSL71^T^.

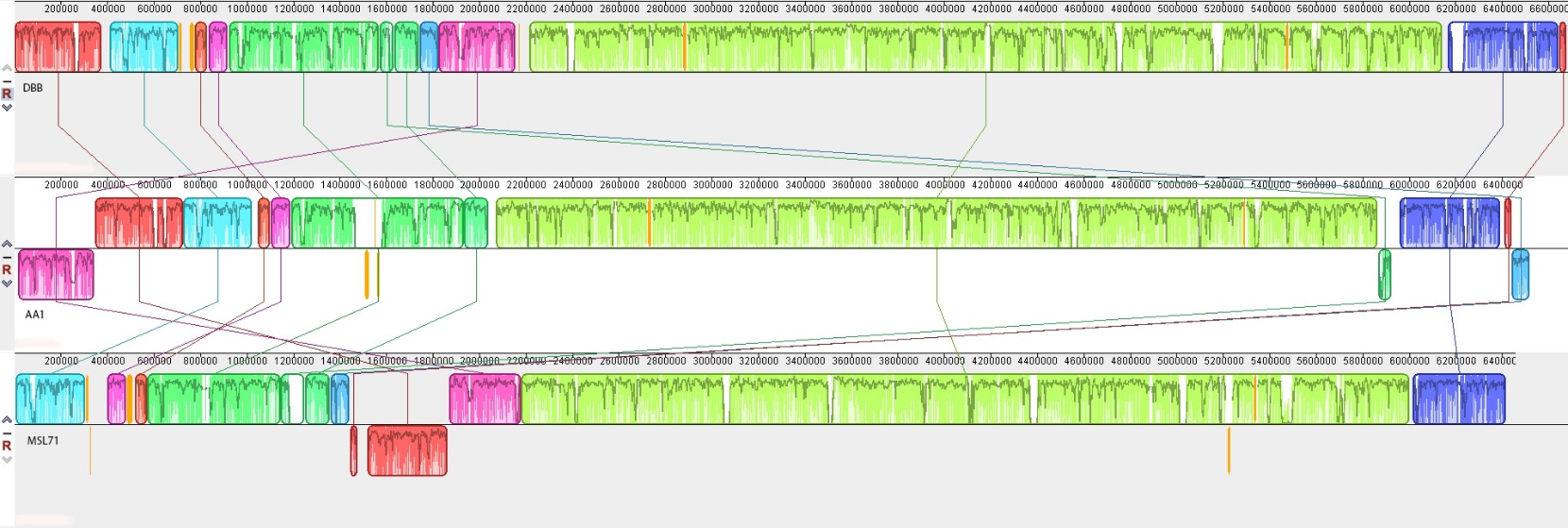

**Fig. S4** Whole genome alignment of *D. spongiiphila* DBB (Top), *D. spongiiphila* AA1^T^ (middle) and *D. butyratoxydans* MSL71^T^ (bottom). The genome of strain DBB was used as reference for global alignment using progressive MAUVE [11]. The locally collinear blocks (LCBs) that were identified in the genomes were outlined in frame. Conserved and highly related regions are coloured, and low-identity unique regions are in white (colorless). LCBs below the mid-line in *D. spongiiphila* AA1^T^ and *D. butyratoxydans* MSL71^T^ are inverted relative to *D. spongiiphila* DBB.

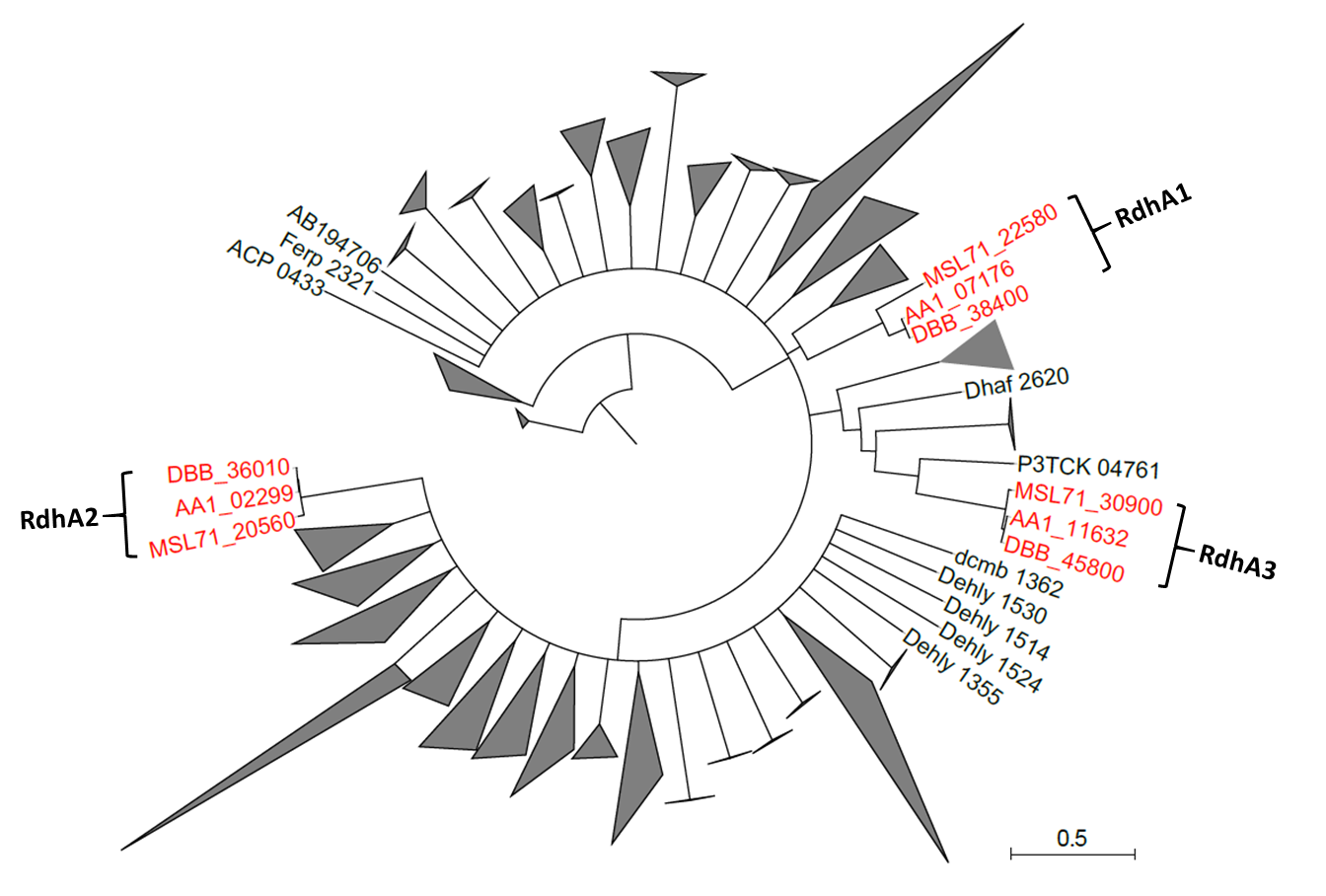

**Fig. S5** Phylogenetic analysis of the RdhAs of *Desulfoluna* strains and 548 RdhAs reported previously [12]. The RdhA sequences were obtained from the public link: <https://drive.google.com/drive/folders/0BwCzK8wzlz8ON1o2Z3FTbHFPYXc>. The multiple sequence alignment was processed using Geneious software with the MAFFT algorithm, and the phylogenetic tree was constructed using the same software with default settings. Further polishing of the phylogenetic tree was performed on the Interactive Tree of Life web browser (<http://itol.embl.de/>) [13]. The RdhAs of *Desulfoluna* strains are shown in red font.

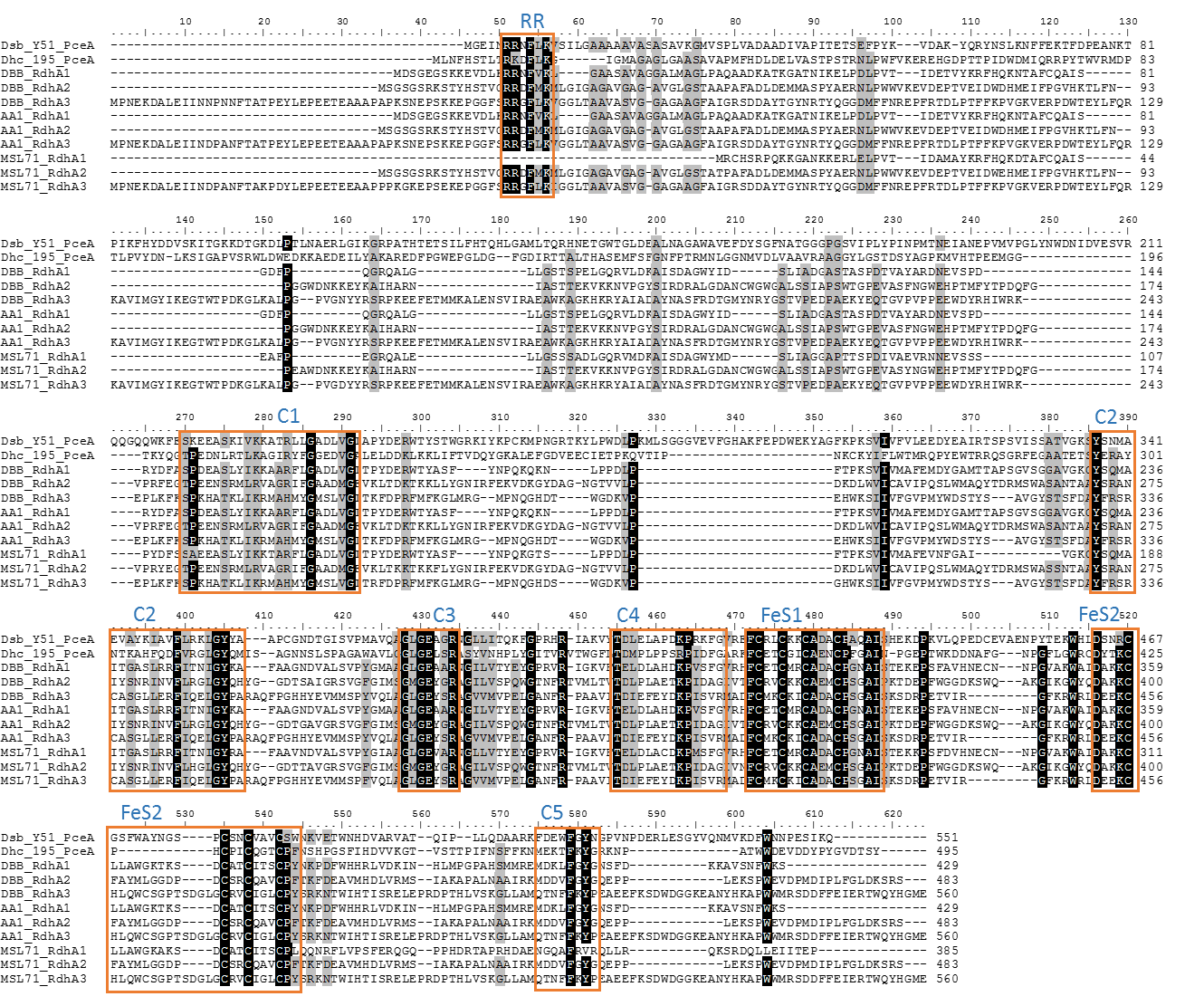

**Fig. S6** Multiple-sequence alignment of the RdhAs from *D. spongiiphila* DBB, *D. spongiiphila* AA1^T^ and *D. butyratoxydans* MSL71^T^ and two functionally characterized RdhAs from *Desulfitobacterium hafniense* Y51, and *Dehalococcoides mccartyi* strain 195. The conserved sequence motifs (RR, C1−C5b FeS1, and FeS2) are enclosed within orange boxes. The selected RdhAs (GenBank accession number or locus number) and corresponding bacteria are: Dsb_Y51_PceA: *D. hafniense* Y51, BAC00915. Dhc_195_PceA: *D. mccartyi* 195, Q3Z9N3. DBB_3755: *D. spongiiphila* DBB. DBB_3984: *D. spongiiphila* DBB. DBB_4749: *D. spongiiphila* DBB. AA1_02299: *D. spongiiphila* AA1^T^. AA1_07176: *D. spongiiphila* AA1^T^. DBB_11632: *D. spongiiphila* AA1^T^. MSL71_1800: *D. butyratoxydans* MSL71^T^. MSL71_2003: *D. butyratoxydans* MSL71^T^. MSL71_4258: *D. butyratoxydans* MSL71^T^

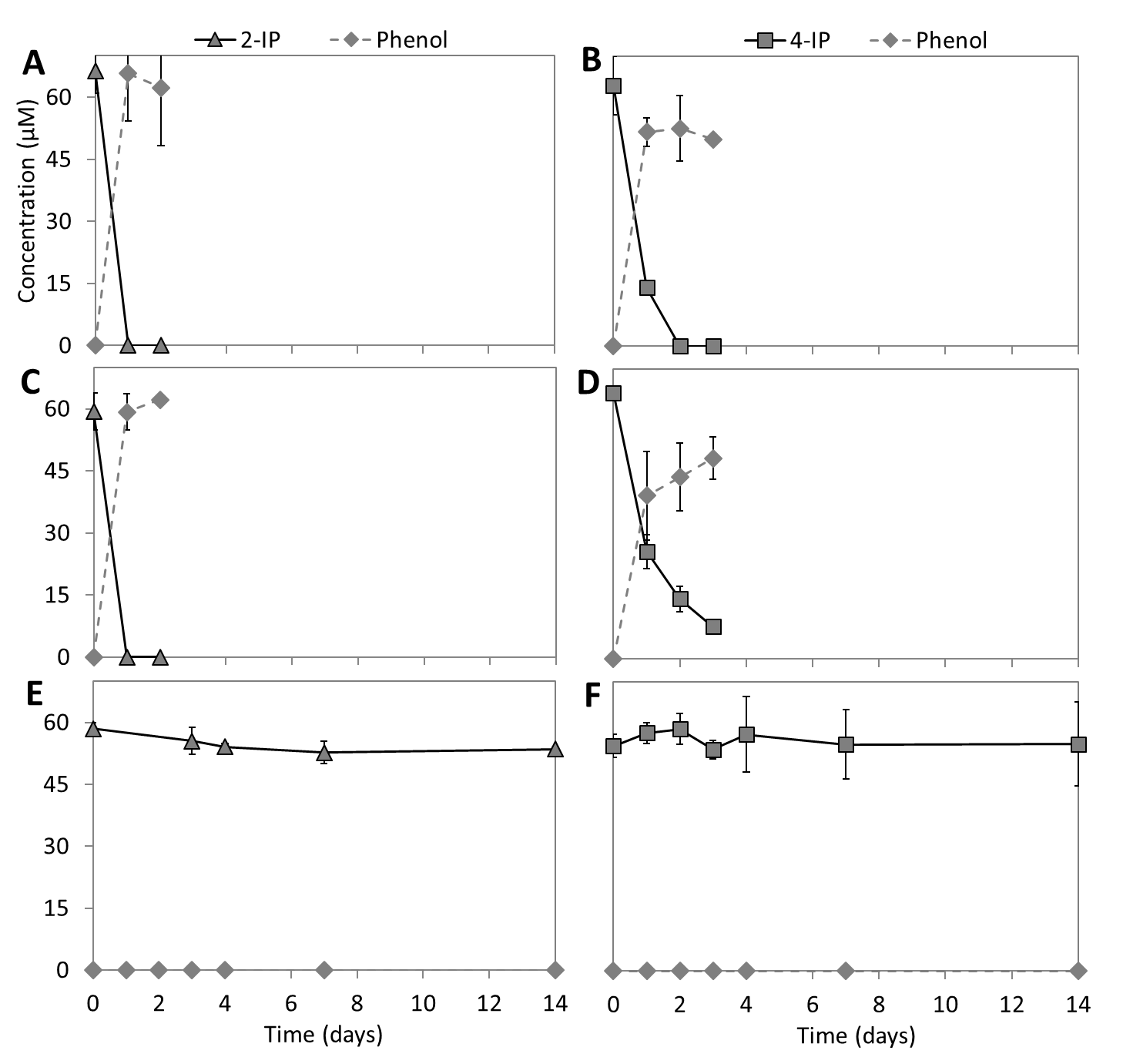

**Fig. S7** Deiodination of 2-IP and 4-IP by *D. spongiiphila* DBB (A, B), *D. spongiiphila* AA1^T^ (C, D) and *D. butyratoxydans* MSL71^T^ (E, F) with lactate (5 mM) as the electron donor. Points and error bars represent the average and standard deviation of samples taken from duplicate cultures.

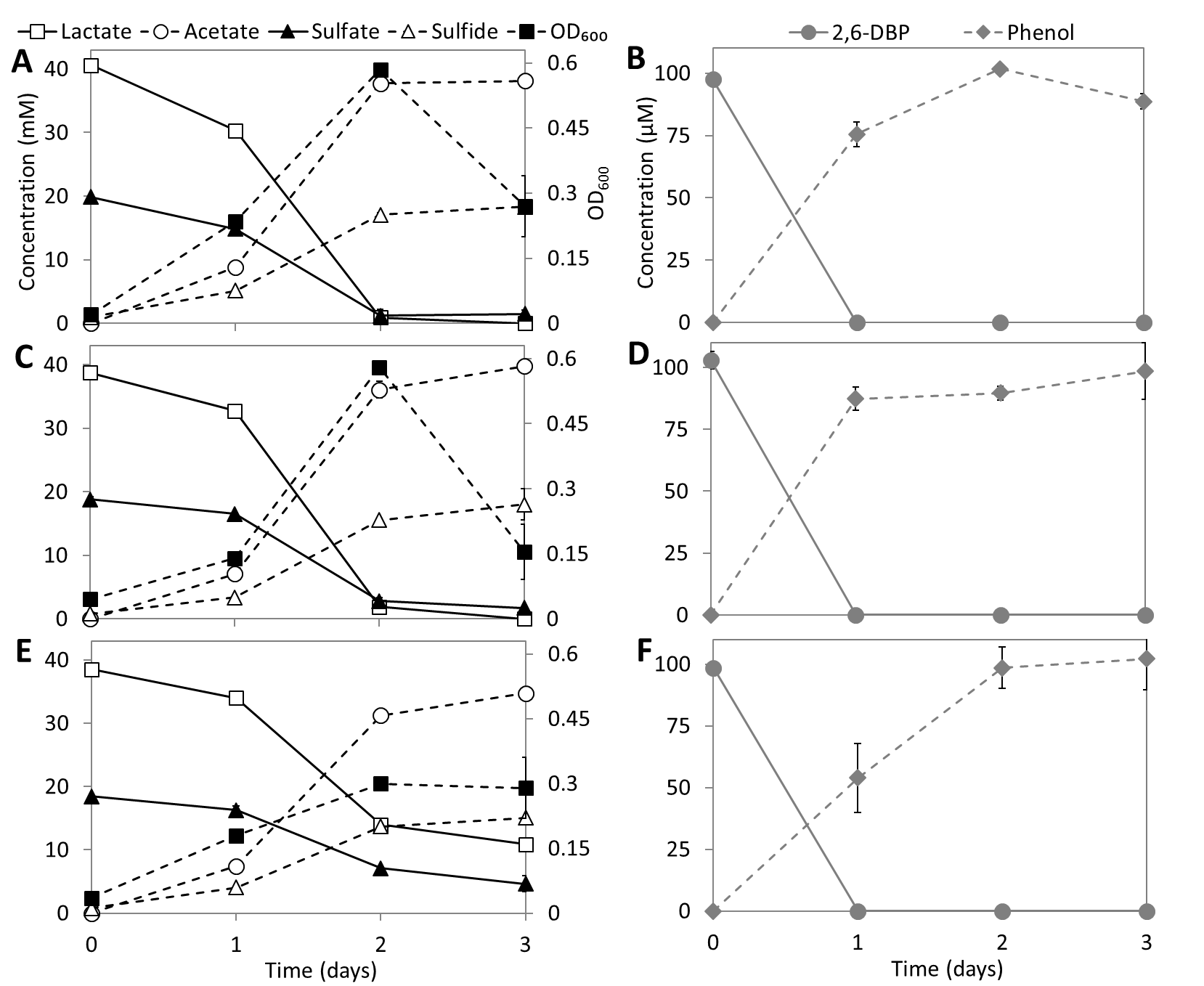

**Fig. S8** Concurrent debromination of 2,6-DBP (100 µM) and sulfate (20 mM) reduction by *D. spongiiphila* DBB (A, B), *D. spongiiphila* AA1^T^ (C, D) and *D. butyratoxydans* MSL71^T^ (E, F) with lactate (40 mM) as the electron donor. Points and error bars represent the average and standard deviation of samples taken from duplicate cultures.

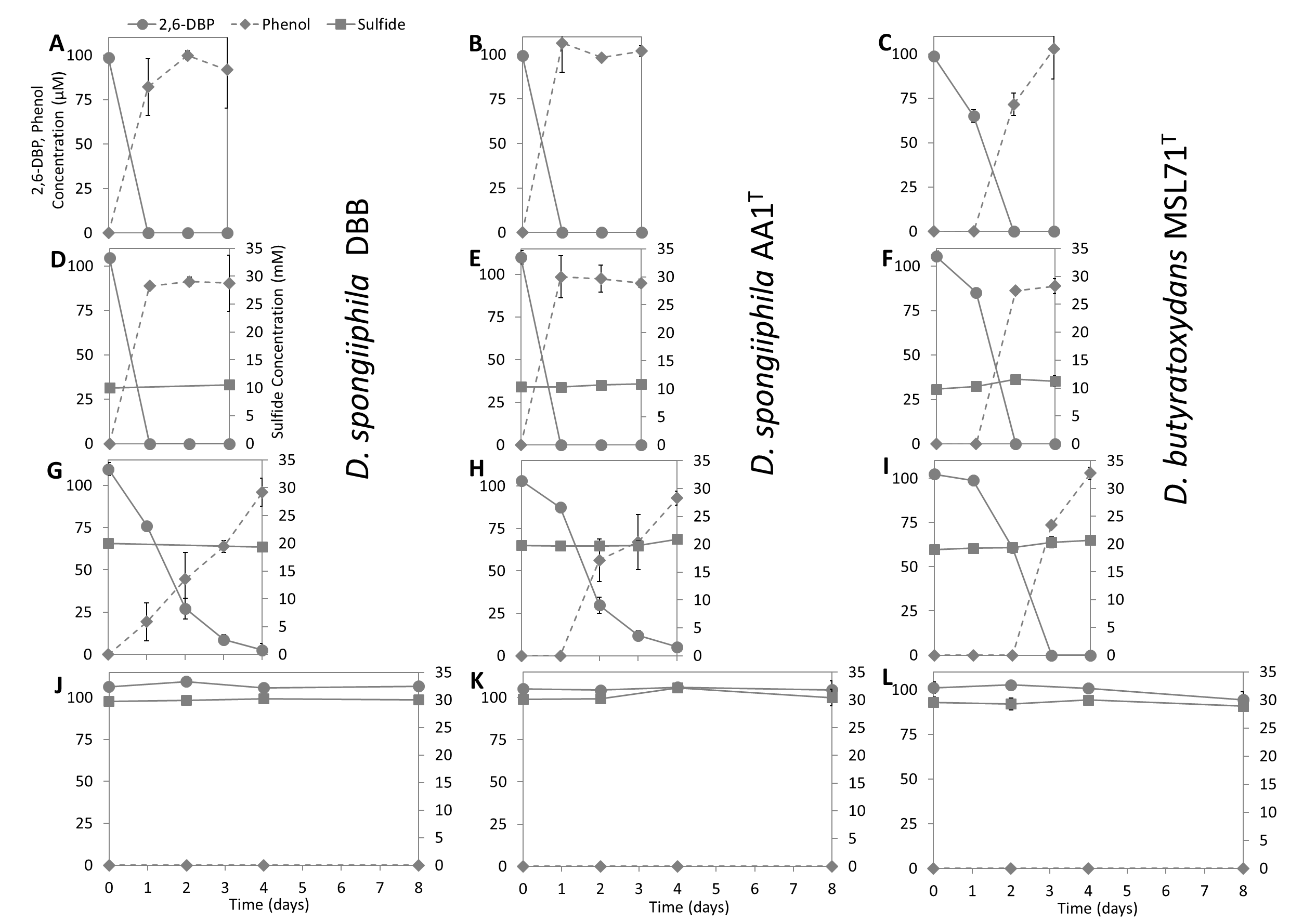

**Fig. S9** Debromination of 2,6-DBP by *D. spongiiphila* DBB (A, D, G, J), *D. spongiiphila* AA1^T^ (B, E, H, K) and *D. butyratoxydans* MSL71^T^ (C, F, I, L) in presence of 1 (A, B, C), 10 (D, E, F), 20 (G, H, I) and 30 mM (J, K, L) sulfide. Points and error bars represent the average and standard deviation of samples taken from duplicate cultures.

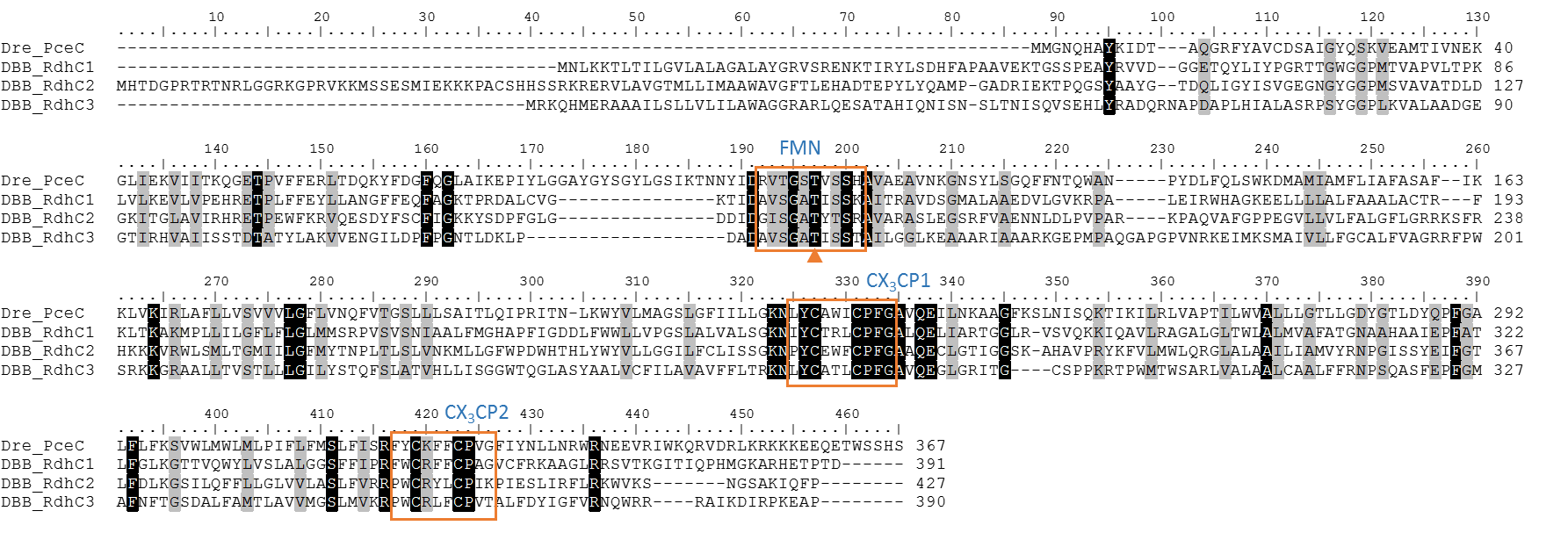

**Fig. S10** Multiple-sequence alignment of the RdhCs of *Desulfoluna spongiiphila* DBB and *Dehalobacter restrictus* (Dre) (GenBank accession number: CAG70347.1). The conserved FMN binding motifs and two CX_3_CP motifs are enclosed within orange boxes. The conserved threonine residue predicted to covalently bind to FMN is indicated with an orange triangle.

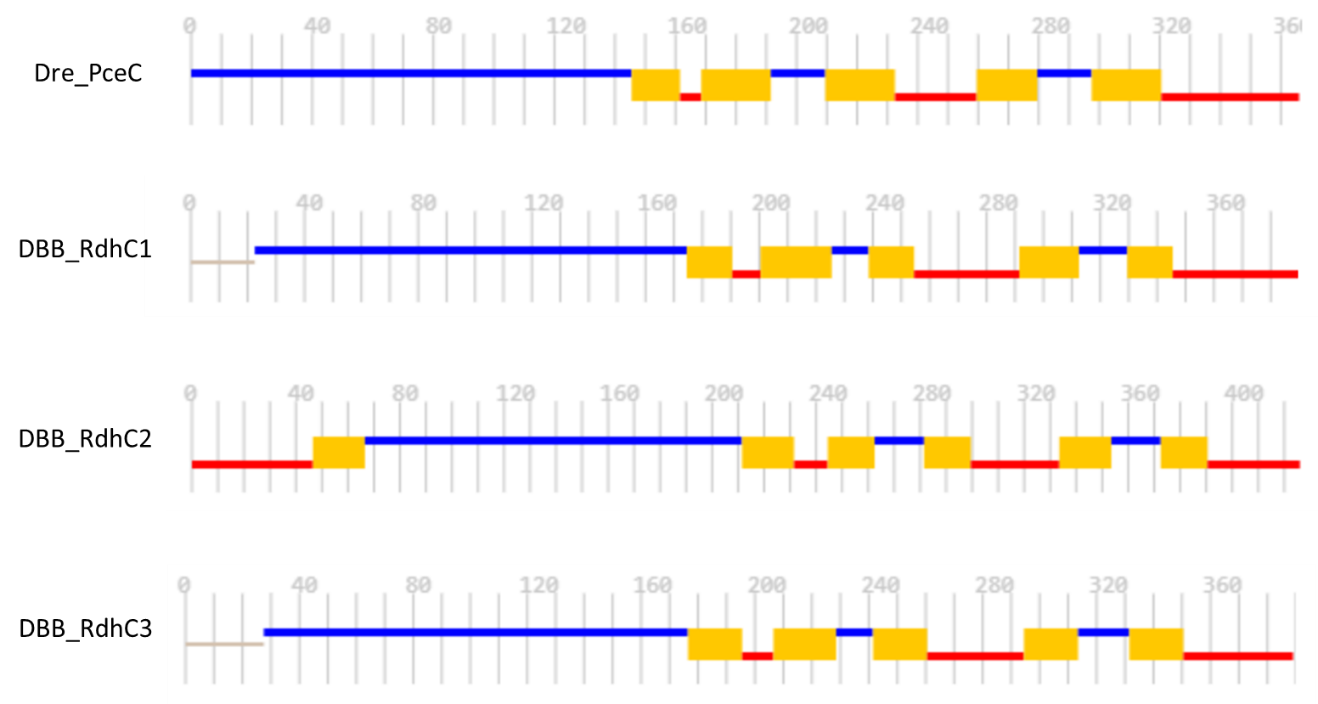

**Fig. S11** Topology analysis and comparison of the RdhCs in *D. spongiiphila* DBB and PceC of *Dehalobacter restrictus*. The topology was predicted using CCTOP [14]. Blue lines indicate outside/extra-cytosolic regions. Red lines indicate inside/cytosolic regions. Gray lines indicate regions where topology is not predicted. Yellow rectangles indicate transmembrane regions.

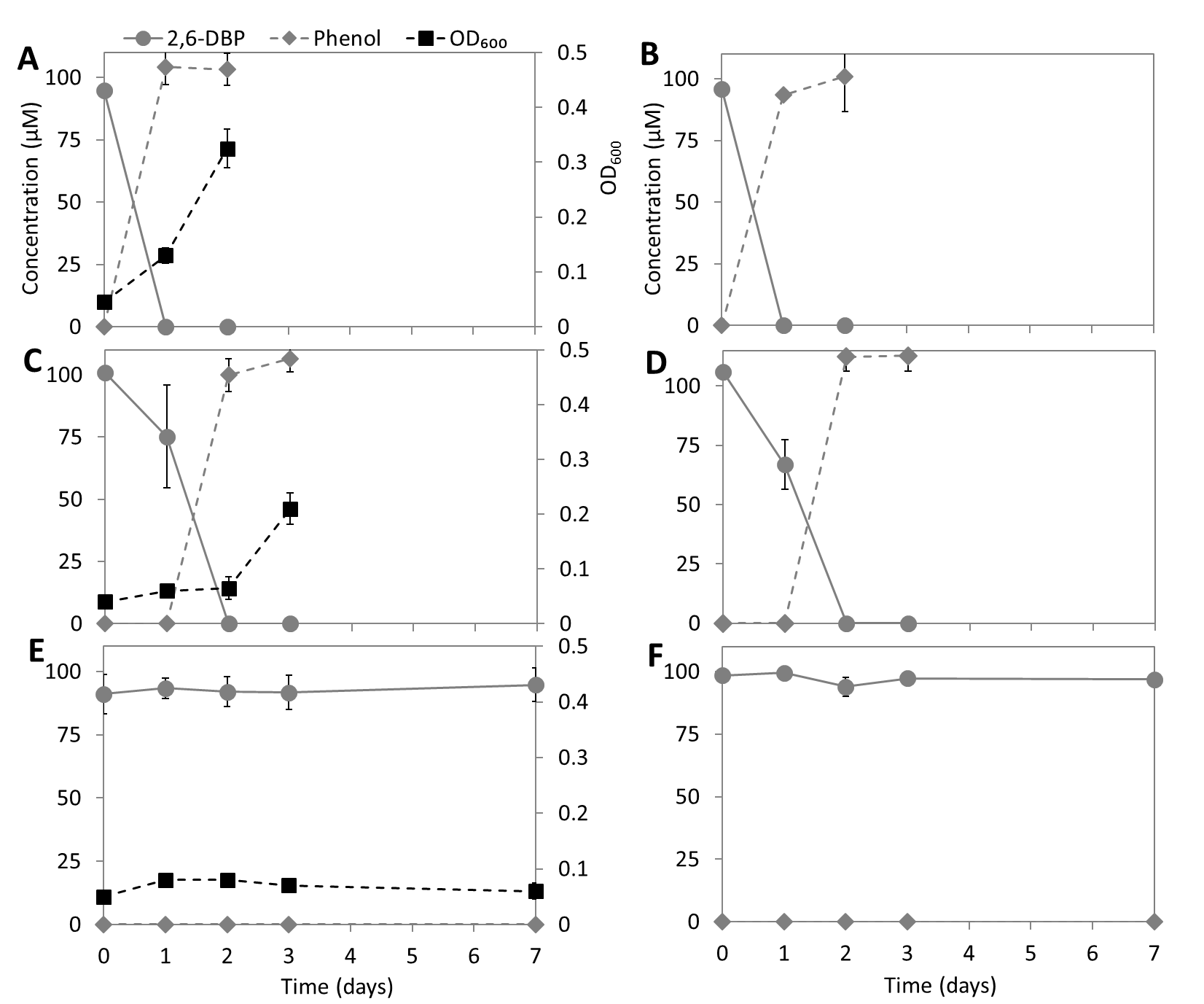

**Fig. S12** 2,6-DBP debromination by *D. spongiiphila* DBB grown in presence (A, C, E) or absence (B, D, F) of sulfate (20 mM) and initial oxygen concentration of 0% (A, B), 2% (C, D), 5% (E, F). Points and error bars represent the average and standard deviation of samples taken from duplicate cultures.

**Supplementary Tables**

Table S1. Primers used in this study.

| Target | Name | Sequence (5´—3´)^a^ | Application | Reference |
| --- | --- | --- | --- | --- |
| Bacteria 16S rRNA | 27F-DegS | GTTYGATYMTGGCTCAG | Miseq | [15] |
|  | 338R–I | GCWGCCTCCCGTAGGAGT | Miseq | [16] |
|  | 338R–II | GCWGCCACCCGTAGGTGT |  |  |
|  | Unitag1 | GAGCCGTAGCCAGTCTGC | Miseq | [17] |
|  | Unitag2 | GCCGTGACCGTGACATCG |  |  |
| Bacteria 16S rRNA | Eub341F | CCTACGGGAGGCAGCAG | qPCR | [18] |
|  | Eub534R | ATTACCGCGGCTGCTGGC |  |  |
| *rdhA1* ^b^ | Rdh1F | ACCGCTACGATTTTGCATCC | qPCR | This study |
|  | Rdh1R | CCATCTCAAAGGCCATGACG |  |  |
| *rdhA*2 ^b^ | Rdh2F | CGTTATTCCGCAGTCGTTGT | qPCR | This study |
|  | Rdh2R | CACTGGGGACTGACAAGGAT |  |  |
| *rdhA*3 ^b^ | Rdh3F | TGGCCGACTTTTGCATGAAA | qPCR | This study |
|  | Rhd3R | AGGTGTTCTTGCGGGAGTAA |  |  |

^a^ M = A or C; R = A or G; W = A or T; Y = C or T

^b^ *rdhA* genes of *D. spongiiphila* DBB

Table S2. Cellular fatty acid composition (%) of different *Desulfoluna* strains

| Fatty acid | *D. spongiiphila* DBB | *D. spongiiphila* AA1^T^ | *D. butyratoxydans* MSL71^T^ |
| --- | --- | --- | --- |
| C12:0 | 0.3 | 0.3 | 0.8 |
| C14:1ω7 | 0.9 | 0.3 | 1.1 |
| C14:1ω5 | 0.2 | 0.1 | 0.2 |
| C14:0 | 9.9 | 6.6 | 11.5 |
| C16:1ω9 | 1.4 | 1.1 | 1.5 |
| C16:1ω7c | 19.4 | 18.9 | 22.4 |
| C16:1ω7tr | 0.3 | 0.3 | 0.3 |
| C16:1ω5 | 0.6 | 0.8 | 0.7 |
| C16:0 | 28.1 | 28.4 | 21.7 |
| C18:1ω9 | 11.6 | 11.8 | 11.3 |
| C18:1ω7 | 18.6 | 22.9 | 19.7 |
| C18:0 | 2.2 | 1.1 | 1.7 |
| β-OH-C12 | 0.4 | 0.5 | 0.4 |
| β-OH-C14 | 6.2 | 7.0 | 6.7 |

Table S3. Selected genomes and their general features that were used for 16S rRNA gene and protein domain based phylogenetic analyses

| Strain ^a^ | Genome size (Mb) | GC (%) | Proteins | GenBank accession number |
| --- | --- | --- | --- | --- |
| *Desulfoluna spongiiphila* DBB | 6.68 | 57.1 | 5301 | xxx |
| *Desulfoluna butyratoxydans* MSL71^T^ | 6.05 | 57.9 | 4186 | xxx |
| *Desulfoluna spongiiphila* AA1^T^ | 6.53 | 57.2 | 5203 | NZ_FMUX01000001.1 |
| *Desulfatibacillum aliphaticivorans* CV2803 | 6.47 | 54.4 | 5264 | NZ_AUCT00000000.1 |
| *Desulfovibrio fructosivorans* JJ | 4.67 | 63.9 | 4046 | NZ_AECZ01000001.1 |
| *Desulfatibacillum alkenivorans* AK-01 | 6.49 | 54.7 | 5277 | NC_011768.1 |
| *Desulfatirhabdium butyrativorans* HB1 | 4.48 | 54.9 | 3852 | NZ_KE386985.1 |
| *Desulfatitalea tepidiphila* S28bF | 5.61 | 56.7 | 4858 | NZ_BCAG01000003.1 |
| *Desulfobacter postgatei* 2ac9 | 3.97 | 47.2 | 3845 | NZ_CM001488.1 |
| *Desulfobacterium autotrophicum* HRM2 | 5.65 | 48.7 | 4835 | NC_012108.1 |
| *Desulfobacterium vacuolatum* DSM 3385 | 5.03 | 46.5 | 4050 | NZ_FWXY01000001.1 |
| *Desulfobacula phenolica* DSM 3384 | 4.87 | 41.4 | 4181 | NZ_FNLL01000001.1 |
| *Desulfobacula toluolica* Tol2 | 5.19 | 41.4 | 4545 | NC_018645.1 |
| *Desulfococcus multivorans* DSM 2059 | 4.42 | 56.8 | 3783 | NZ_CP015381.1 |
| *Desulfococcus oleovorans* Hxd3 | 3.94 | 56.2 | 3325 | NC_009943.1 |
| *Desulfomicrobium baculatum* DSM 4028 | 3.94 | 58.6 | 3395 | NC_013173.1 |
| *Desulfosarcina cetonica* JCM 12296 | 7.09 | 55.7 | 5582 | NZ_BBCC01000001.1 |
| *Desulfotignum phosphitoxidans* DSM 13687 | 4.99 | 51.3 | 4556 | NZ_APJX01000001.1 |
| *Desulfovibrio aespoeensis* Aspo 2 | 3.62 | 62.6 | 3257 | NC_014844.1 |
| *Desulfovibrio alaskensis* G20 | 3.64 | 57.9 | 3270 | NC_007519.1 |
| *Desulfovibrio desulfuricans* ND132 | 3.85 | 65.2 | 3423 | NC_016803.1 |

^a^ Genome information were obtained from GenBank under their respective accession numbers, except *D. spongiiphila* DBB and *D. butyratoxydans* MSL71^T^

Table S4. Abundance of the proteins involved in lactate, sulfate and 1,4-DBB metabolism in cells of strain DBB grown in LS and LSD conditions

|  | Locus tag | LS1 Area | LS2 Area | LS3 Area | LSD1 Area | LSD2 Area | LSD3 Area | log2 fold-change | p-value |
| --- | --- | --- | --- | --- | --- | --- | --- | --- | --- |
| Proteins involved in lactate metabolism | | | | | | | | | |
| Lactate permease | 24890 | 23.64 | 23.48 | 24.38 | 22.96 | 23.17 | 24.12 | 0.42 | 0.40 |
| LdhA-1 | 24880 | 28.72 | 28.66 | 29.09 | 29.09 | 29.01 | 28.59 | 0.08 | 0.73 |
| LdhA-2 | 24970 | 28.03 | 28.08 | 28.37 | 27.89 | 27.85 | 27.88 | -0.28 | 0.05 |
| LdhB-1 | 24870 | 25.39 | 25.97 | 26.61 | 26.50 | 25.84 | 25.93 | 0.10 | 0.81 |
| LdhB-2 | 24960 | 26.35 | 26.69 | 29.20 | 26.84 | 25.89 | 26.12 | -1.13 | 0.29 |
| Por-1 | 310 | 31.58 | 31.55 | 31.62 | 30.70 | 30.64 | 30.79 | -0.87 | 0.00 |
| Por-2 | 24940 | 32.20 | 32.22 | 32.18 | 31.20 | 31.05 | 31.25 | -1.03 | 0.00 |
| Pta | 9370 | 27.45 | 27.31 | 27.91 | 27.97 | 27.56 | 27.57 | 0.15 | 0.55 |
| Ack | 9360 | 28.26 | 28.72 | 28.92 | 29.10 | 28.35 | 28.27 | -0.06 | 0.86 |
| Proteins involved in sulfate metabolism | | | | | | | | | |
| Sulfate permease | 22290 | 22.35 | 22.14 | 22.49 | 21.25 | 21.21 | 22.95 | 0.52 | 0.42 |
| Sat | 23930 | 30.04 | 30.03 | 29.50 | 29.71 | 30.25 | 29.73 | -0.04 | 0.88 |
| ApsBA | 23880 | 26.33 | 26.34 | 25.92 | 26.22 | 25.94 | 26.11 | 0.11 | 0.54 |
|  | 23890 | 29.22 | 28.85 | 28.57 | 29.21 | 28.63 | 29.02 | -0.08 | 0.78 |
| QmoABC | 23900 | 24.36 | 23.64 | 24.40 | 23.92 | 23.45 | 24.16 | 0.29 | 0.42 |
|  | 23910 | 24.88 | 24.32 | 24.36 | 24.18 | 23.44 | 24.48 | 0.48 | 0.25 |
|  | 23920 | 26.22 | 26.24 | 26.34 | 26.10 | 26.30 | 26.72 | -0.11 | 0.59 |
| DsrC | 370 | 31.27 | 31.08 | 31.57 | 31.58 | 30.91 | 31.38 | -0.02 | 0.95 |
| DsrABD | 25620 | 32.60 | 32.22 | 32.75 | 32.50 | 32.82 | 32.47 | 0.08 | 0.71 |
|  | 25630 | 32.66 | 32.45 | 32.50 | 32.32 | 32.18 | 32.31 | -0.26 | 0.03 |
|  | 25640 | 29.06 | 28.67 | 29.24 | 29.11 | 29.04 | 28.98 | 0.06 | 0.77 |
| DsrMKJOP | 27290 | 22.46 | 21.95 | 23.37 | 22.84 | 22.29 | 22.87 | -0.08 | 0.87 |
|  | 27300 | 24.46 | 23.24 | 22.96 | 22.72 | 23.43 | 24.32 | 0.07 | 0.92 |
|  | 27310 | ND | ND | ND | ND | ND | ND | - | - |
|  | 27320 | 22.28 | 19.51 | 22.20 | 22.90 | 22.59 | 22.90 | -1.46 | 0.18 |
|  | 27330 | 22.72 | 23.09 | 23.97 | 22.80 | 22.52 | 23.81 | 0.22 | 0.70 |
| Electron transport proteins | | | | | | | | | |
| FixABC | 25970 | 30.04 | 29.82 | 30.06 | 30.29 | 30.65 | 30.11 | 0.37 | 0.09 |
|  | 25980 | 29.59 | 29.14 | 29.98 | 29.82 | 30.32 | 29.85 | 0.42 | 0.22 |
|  | 25990 | 23.32 | 22.36 | 23.48 | 21.62 | 20.86 | 23.02 | 1.22 | 0.17 |
| Flavodoxin | 37290 | 33.29 | 33.59 | 31.86 | 32.73 | 33.16 | 33.91 | 0.35 | 0.61 |
| QrcABCD | 34140 | ND | ND | ND | ND | ND | ND | - | - |
|  | 34150 | 20.66 | 21.51 | 21.22 | 22.42 | 21.33 | 22.31 | -0.89 | 0.11 |
|  | 34160 | 22.10 | 22.60 | 22.56 | 22.18 | 22.28 | 22.82 | -0.01 | 0.97 |
|  | 34170 | ND | ND | ND | ND | ND | ND | - | - |
| Reductive dehalogenase | | | | | | | | | |
| RdhA1 | 38400 | ND | ND | ND | 26.62 | 25.99 | 25.39 | - | - |
| Corrinoid biosynthesis proteins | | | | | | | | | |
| GltX | 24080 | 25.00 | 24.89 | 24.99 | 24.83 | 24.45 | 24.66 | -0.31 | 0.05 |
| HemL | 7500 | 26.66 | 27.12 | 26.78 | 26.39 | 26.04 | 24.88 | -1.08 | 0.08 |
| HemB | 44050 | 26.25 | 26.89 | 26.29 | 25.94 | 25.78 | 26.45 | -0.42 | 0.22 |
| HemC | 18940 | 27.66 | 27.70 | 27.57 | 27.16 | 27.31 | 27.64 | -0.27 | 0.14 |
| HemD | 18950 | 27.94 | 27.84 | 27.70 | 27.71 | 27.83 | 26.97 | -0.32 | 0.31 |
| CysG | 26600 | 24.85 | 25.24 | 25.21 | 25.04 | 25.02 | 25.05 | -0.06 | 0.63 |
| CbiK | 3730 | 22.33 | 23.47 | 22.80 | 21.70 | 22.66 | 22.96 | -0.43 | 0.44 |
| CbiL | 3790 | 24.06 | 24.62 | 23.45 | 24.55 | 23.59 | 24.36 | 0.12 | 0.79 |
| CbiH | 3850 | 23.81 | 24.54 | 24.96 | 25.56 | 25.70 | 26.10 | 1.35 | 0.02 |

The full name of each protein can be found in Figure 6 and Table S5

ND: not detected

Table S5. Corrinoid biosynthesis pathways and corresponding genes and functions in *Desulfoluna* strains

| Biosynthetic pathway | Gene ^a^ | DBB ^b^ | AA1^T c^ | MSL71^T b^ | Function in corrinoid biosynthesis |
| --- | --- | --- | --- | --- | --- |
| Glutamate |  |  |  |  |  |
| ↓ | *gltX* | **24080** | 11166 | 15970 | Glutaminyl-trna synthetase |
| ↓ | *hemA* | 26620 | 12922 | 13190 | Glutamyl-trna reductase |
| ↓ | *hemL* | **7500** | 10673 | 46530 | Glutamate-1-semialdehyde 2,1-aminomutase |
| ↓ | *hemB* | **44050** | 13043 | 37790 | Porphobilinogen synthase |
| ↓ | *hemC* | **18940** | 103136 | 7120 | Hydroxymethylbilane synthase |
| ↓ | *hemD* | **18950** | 103137 | 7130 | Uroporphyrinogen-III synthase |
| Uroporpyhrinogen III |  |  |  |  |  |
| ↓ | *cysG* | **26600** | 12920 | 13210 | Uroporphyrin-III C-methyltransferase |
| Precorrin-2 |  |  |  |  |  |
| ↓ | *cbiK* | **3730** | 12810 | 49290 | Sirohydrochlorin cobaltochelatase |
| Co(II)- precorrin-2 |  |  |  |  |  |
| ↓ | *cbiL* | **3790** | 12816 | 49350 | Precorrin-2/cobalt-factor-2 C20-methyltransferase |
| Co(II)- precorrin-3 |  |  |  |  |  |
| ↓ | *cbiH* | **3850** | 12822 | 49410 | Precorrin-3B C17-methyltransferase |
| Co(II)-precorrin-4 |  |  |  |  |  |
| ↓ | *cbiF* | 3830 | 12820 | 49390 | Precorrin-4/cobalt-precorrin-4 C11-methyltransferase |
| Co(II)-precorrin-5A |  |  |  |  |  |
| ↓ | *cbiG* | 3840 | 12821 | 49400 | Cobalt-precorrin 5A hydrolase |
| Co(II)-precorrin-5B |  |  |  |  |  |
| ↓ | *cbiD* | 3810 | 12818 | 49370 | Cobalt-precorrin-5B (C1)-methyltransferase |
| Co(II)-precorrin-6A |  |  |  |  |  |
| ↓ | *cbiJ* ^d^ |  |  |  | Precorrin-6A/cobalt-precorrin-6A reductase |
| Co(II)-precorrin-6B |  |  |  |  |  |
| ↓ | *cbiET* | 3820 | 12819 | 49380 | Cobalt-precorrin-6B (C15)-methyltransferase |
| Co(II)-precorrin-7,8 |  |  |  |  |  |
| ↓ | *cbiC* | 3780 | 12815 | 49340 | Precorrin-8X/cobalt-precorrin-8 methylmutase |
| Cobyrinic acid |  |  |  |  |  |
| ↓ | *cbiA* | 3770 | 12814 | 49330 | Cobyrinic acid *a,c*-diamide synthase |
| Cob(II)yrinic acid *a,c*-diamide |  |  |  |  |  |
| ↓ |  |  |  |  |  |
| Cob(I)yrinic acid *a,c*-diamide |  |  |  |  |  |
| ↓ | *cobA* | 3860 | 12823 | 49420 | Cob(I)alamin adenosyltransferase |
| Ado-cob(I)yrinic acid *a,c*-diamide |  |  |  |  |  |
| ↓ | *cbiP* | 3870 | 12824 | 49430 | Adenosylcobyric acid synthase |
| Adenosyl cobyrinate *a,c*-hexaamide |  |  |  |  |  |
| ↓ | *cbiB* | 3920 | 12829 | 49480 | Adenosylcobinamide-phosphate synthase |
| Ado-cobinamide |  |  |  |  |  |
| ↓ | *cobU* | 3880 | 12825 | 49440 | Adenosylcobinamide-phosphate guanylyltransferase |
| Ado-cobinamide-GDP |  |  |  |  |  |
| ↓ | *cobS* | 3890 | 12826 | 49450 | Cobalamin synthase |
| Cobalamin |  |  |  |  |  |

^a^ Gene nomenclature for the anaerobic corrinoid biosynthesis pathway was as previously published [19]

^b^ Gene Locus numbers are according the genome sequences of strains DBB and MSL71^T^ sequenced in this study. The numbers shown in bold are proteins detected in the proteome of strain DBB

^c^ Gene Locus numbers are according the genome sequence of stain AA1^T^ in GenBank

^d^ Genes that were not found in the *Desulfoluna* genomes

Table S6. Sulfur metabolism pathways and corresponding genes in *Desulfoluna* strains

| Metabolic pathway | Enzyme | DBB ^a^ | AA1^T b^ | MSL71^T a^ |
| --- | --- | --- | --- | --- |
|  | Sulfate permease | 13340, **22290**, 24700, 47100 | 12364, 11742, 111127, 104190 | 27680, 17610, 15410, 29700 |
| Tetrathionate |  |  |  |  |
| ↓ | Tetrathionate reductase | 32070, 32080 | 11084, 11085 | ND |
| Thiosulfate |  |  |  |  |
| ↓ | Molybdopterin oxidoreductase  or  Rhodanese-like protein^c^ | 8600—8620    9670, 10640 | 106177—106179  10968, 10571 | 41260—41280  51880, 25130 |
| Sulfite |  |  |  |  |
| Sulfate |  |  |  |  |
| ↓ | Sulfate adenylyltransferase | **23930** | 11149 | 16120 |
| Adenylyl sulfate (APS) |  |  |  |  |
| ↓ | APS reductase α subunit | **23890** | 11145 | 16160 |
|  | APS reductase β subunit | **23880** | 11144 | 16170 |
| Sulfite |  |  |  |  |
| ↓ | Dissimilatory sulfite reductase α subunit | **25620** | 12530 | 14160 |
|  | Dissimilatory sulfite reductase β subunit | **25630** | 12529 | 14150 |
|  | Dissimilatory sulfite reductase D | **25640** | 12528 | 14140 |
| Sulfide |  |  |  |  |

^a^ Locus tag numbers are according to the genome of strains DBB and MSL71^T^ sequenced in this study. Numbers shown in bold are proteins detected in the proteome of strain DBB

^b^ Locus tag numbers are according to the genome of stain AA1^T^ (NZ_FMUX01000001.1)

^c^ Putative function as thiosulfate reductase

ND: Not detected

Table S7. Enzymes involved in oxygen reduction and ROS detoxification in *Desulfoluna* strains

| Enzyme | DBB ^a^ | AA1^T b^ | MSL71^T a^ |
| --- | --- | --- | --- |
| Rubredoxin–oxygen oxidoreductase | 16300 | 101503 | 5010 |
| Cytochrome *c* oxidase | 43200–43500 | 11283–11286 | 36840–36870 |
| Cytochrome *bd* oxygen reductase | 15890–15900 | 101463–101464 | 4610–4620 |
| Superoxide dismutase | 15650 | 101439 | 4380 |
| Superoxide reductase | 40040 | 1079 | 24220 |
| Rubrerythrin | 34920 | 102199 | 19140 |
| Thiol peroxidase | 30870 | 114122 | 33350 |

^a^ Locus tag numbers are according to the genomes of strains DBB and MSL71^T^ sequenced in this study

^b^ Locus tag numbers are according to the genome of stain AA1^T^ (NZ_FMUX01000001.1)
